# MIP-MAP: High Throughput Mapping of *Caenorhabditis elegans* Temperature Sensitive Mutants via Molecular Inversion Probes

**DOI:** 10.1101/150862

**Authors:** CA Mok, V Au, OA Thompson, ML Edgley, L Gevirtzman, J Yochem, J Lowry, N Memar, M Wallenfang, D Rasoloson, B Bowerman, R Schnabel, G Seydoux, DG Moerman, RH Waterston

## Abstract

Temperature sensitive (TS) alleles are important tools for the genetic and functional analysis of essential genes in many model organisms. While isolating TS alleles is not difficult, determining the TS-conferring mutation can be problematic. Even with whole-genome sequencing (WGS) data there is a paucity of predictive methods for identifying TS alleles from DNA sequence alone. We assembled 173 TS lethal mutants of *Caenorhabditis elegans* and used WGS to identify several hundred mutations per strain. We leveraged single molecule molecular inversion probes (MIPs) to sequence variant sites at high depth in the cross-progeny of TS mutants and a mapping strain with identified sequence variants but no apparent phenotypic differences from the reference N2 strain. By sampling for variants at ~1Mb intervals across the genome we genetically mapped mutant alleles at a resolution comparable to current standards in a process we call MIP-MAP. The MIP-MAP protocol, however, permits high-throughput sequencing of multiple TS mutation mapping libraries at less than 200K reads per library. Using MIP-MAP on a subset of TS mutants, via a competitive selection assay and standard recombinant mutant selection, we defined TS-associated intervals of 3Mb or less. Our results suggest this collection of strains contains a diverse library of TS alleles for genes involved in development and reproduction. MIP-MAP is a robust method to genetically map mutations in both viable and essential genes. The MIPs protocol should allow high-throughput tracking of genetic variants in any mixed population.

## Introduction

The Million Mutation Project, a collection of genomic data for thousands of mutagenized *C. elegans* strains (Thompson *et al.* 2013), has provided a dense library of mutant alleles with which to study gene function and has been widely used since its release. However, given that such strains generally contain only homozygous viable mutations, they largely exclude strongly deleterious alleles of essential genes. Such genes encompass a range of classes that include roles in cell division, development and fertility – key components to all multicellular organisms. Temperature-sensitive (TS) lethal alleles can facilitate the genetic analysis of essential genes through the conditional modulation of their function without the complications that balancer chromosomes can introduce when present in non-conditional lethal strains (Golden *et al.* 2000; O’Rourke *et al.* 2011a; Lowry *et al.* 2015). There are several reported screens for TS lethal alleles in *C. elegans* but to date, there are only a small portion of genes with TS alleles identified (Zonies *et al.* 2010; Ehmke *et al.* 2014; Lowry *et al.* 2015). Generating a comprehensive library of mutant strains with conditional lethal phenotypes has the potential to expand our knowledge of essential genes, their required levels of expression, the timing of their function(s), and the details of their protein and domain structure. Furthermore, such a catalog could have broad implications for elucidating the life cycle of *C. elegans* and organisms beyond this model. Such an endeavour, however, requires the mapping and characterization of mutant alleles in a systematic and high-throughput manner.

Chemical mutagenesis methods to generate TS alleles typically result in up to several hundred mutations per strain (Sarin *et al.* 2008; Flibotte *et al.* 2010; Thompson *et al.* 2013). Determining which of these molecular alterations is responsible for the TS mutant phenotype is especially challenging when dealing with partially penetrant or leaky alleles. Many mutations can initially be prioritized as candidates because they alter the coding potential of a gene, but even after such filtering, more than 1/5 of the collection’s SNVs remain. In contrast to identifying constitutive loss-of-function alleles, such as deletions, frame-shifts and nonsense mutations, there is, as yet, no universal predictive method for recognizing which coding or perhaps noncoding molecular change is most likely to give rise to a temperature sensitive phenotype (Perry *et al.* 1994; Rogalski *et al.* 1995; Harfe *et al.* 1998; Poultney *et al.* 2011). When feasible, mutants are initially outcrossed or backcrossed to remove extraneous mutations originating from the mutagenesis process. Otherwise, these additional mutations may, to some degree, affect development or other essential pathways, thus obfuscating the process of allele characterization.

Genetic mapping is a general method for identifying the causative mutation in such strains. Mapping with classic visible genetic markers in *C. elegans* can be a laborious, iterative process. Single nucleotide variants between strains, such as wildtype N2 and the Hawaiian strain CB4856, act as molecular genetic markers that allow the parallel mapping of multiple sites across the genome in a single cross. The so-called Snip-SNP assay is a popular method that exploits a subset of Hawaiian genome SNVs that alter recognition sites of the restriction enzyme DraI (Davis *et al.* 2005). However, the Hawaiian genome is proven to harbor various alleles (De Bono and Bargmann 1998; Seidel *et al.* 2008, 2011; Andersen *et al.* 2014) that can negatively alter the representation of segregant populations; CB4856 also has phenotypes of its own that may interfere with the scoring of some behavioral phenotypes (Wicks *et al.* 2001). Despite these issues, the research community continues to leverage CB4856 as a mapping strain to develop new methods of mapping complex mutations (Doitsidou *et al.* 2010; O’Rourke *et al.* 2011b; Minevich *et al.* 2012; Smith *et al.* 2016).

More recently, approaches and tools have been developed that allow the simultaneous measurement of SNV frequency in a cross-population. Included amongst these are a combined-step WGS and SNP analysis complemented by analysis via the CloudMap system as well as restriction site associated DNA polymorphism mapping (Doitsidou *et al.* 2010; O’Rourke *et al.* 2011b; Minevich *et al.* 2012). At their core, these mapping methods still rely on the basic principles of bulk segregant analysis except that they now benefit from using massively parallel sequencing to examine molecular markers across the genome in a single sequencing library. As currently implemented these mapping methods use whole genome sequencing (WGS) to a depth of 18-40X or more on populations generated from 20-50 F2 mutant phenotype animals (Doitsidou *et al.* 2010; Wang *et al.* 2014; Jaramillo-Lambert *et al.* 2015; Lowry *et al.* 2015) to capture data on both the recombinant genome landscape and the associated mutant allele. On a large scale, mutant mapping strains is a labor-intensive exercise. The large amount of sequence generation, the use of the Hawaiian strain - with its limitations - and the issues with scaling to high-throughput suggest there is room for improvement. In particular, molecular inversion probes (Turner *et al.* 2009) (MIPs), allow high-throughput, target-based genome amplification and sequencing. A single MIP targets a region via a pair of complementary annealing arms. Once annealed, the genomic sequence between arms, also known as the gap-fill sequence, is copied by a DNA polymerase and the entire single-stranded probe is circularized by ligase. Each circularized probe represents a unique strand of DNA that can now be linearized and sequenced. MIPs have proven very useful in targeted exome sequencing libraries (Mamanova *et al.* 2010; Kiezun *et al.* 2013) and in screening samples for subpopulations of variants. Recently, Hiatt et al., introduced single molecule molecular inversion probes (smMIPs) containing the addition of a randomized molecular barcode to differentiate between low-frequency variants and those contributed by sequencing or amplification artifacts in genomically diverse samples (Hiatt *et al.* 2013).

In complement to the Million Mutation Project collection we have sequenced a group of 173 previously uncharacterized mutants obtained from multiple screens for TS lethality, which reveals a rich catalog of genomic variants. To identify the specific mutation(s) underlying these TS phenotypes, we developed an alternative strategy to whole genome sequencing genetic mapping that exploits MIPs to specifically assay only regions of the genome containing SNVs of interest. To avoid the pitfalls associated with the Hawaiian strain we identified a polymorphic mapping strain from the Million Mutation Project that resembles N2 in movement and growth but has fewer than 300 SNVS across the genome. In pilot experiments, we demonstrate the utility of our MIP-based method (MIP-MAP) in generating high-resolution genomic interval maps for *C. elegans* mutant alleles. We further developed a competitive fitness approach to map the associated genomic intervals for a subset of our TS lethal mutants. In crosses with our mapping strain we use non-permissive temperatures to select against TS lethal homozygotes over several generations to identify candidate intervals and query our sequencing data to produce a list of candidate mutations. Our studies identify TS alleles for a variety of novel genes not previously classified as essential to development or reproduction, suggesting our collection can be a useful resource for studying essential genes.

## Materials and Methods

### TS strain isolation and sequencing

Set 1 strains originated from the Bowerman lab and were isolated from an ENU mutagenesis protocol (Kemphues *et al.* 1988b) in EU1700, a *lin-2(e1309)* background with an integrated *pie-1*-driven series of GFP fusions to beta-tubulin, histone 2B and a PH domain. Mutagenized animals were screened for conditional TS embryonic lethality by shifting F2 animals as L4s from 15°C to 26°C for 24-36 hours and identifying those with an accumulation of dead embryos. These F2 animals were then shifted back to 15°C with the intention of isolating any strains that managed to recover and produce some progeny. Recovered strains were then grown at 15°C for a few generations before rescreening at 26°C to confirm for TS lethality.

Set 2 strains originated from the Schnabel lab and were isolated from EMS mutagenesis (Brenner 1974; Ehmke *et al.* 2014) in an wildtype (N2) background. After 8 days of growth at 15°C, animals were singled to 96-well plates and visually screened for TS lethality at 25°C. Positive wells were regrouped to new 96-well plates twice before testing a second time at 25°C. Second-round positive strains were plated on individual NGM plates before confirming TS lethality a third time at the non-permissive temperature followed by detailed phenotype analysis.

Set 3 strains originated from the Seydoux lab and were isolated from EMS mutagenesis (Kemphues *et al.* 1988b; Golden *et al.* 2000)in a JH50 background (*him-3(e1147)IV; lin-2(e1309) axIS36 X*). F2 animals were screened by bleach synchronization of F1 gravid adults followed by upshift at L4 from 15°C to 25°C for 20 hours and then down to 15°C for an additional 20 hours. F2 animals accumulating dead embryos were singled to NGM plates at 15°C and examined 3 days later for F3 progeny, indicating the presence of a maternal effect embryonic TS lethal mutation.

SNV mutations for all 173 strains were sequenced and analyzed using the same methods and custom pipelines described previously for the Million Mutation Project (Thompson *et al.* 2013). SNV calls for each strain can be found in Supplemental Data SD2.

### MIP-MAP Design

The smMIP design for these experiments was altered from those presented in Hyatt et al. 2013, by placing the 12bp molecular barcode directly on the 3’ end of the first read primer sequence. This alteration allows MIP sequencing to proceed in a single-end 50bp read that includes the molecular barcode, ligation arm sequence, and 18bp of gap-fill sequence. MIP annealing arms were designed using a custom R script that generated multiple combinations of lengths and locations for the ligation and extension arms. Ligation arm locations were limited to a maximum distance of 18bp either up- or down-stream from the SNV of interest. From these possible variations, the optimal MIP sequences were then chosen based on criteria as described elsewhere (Turner *et al.* 2009; O’Roak *et al.* 2012). MIP sequence information for VC20019 probes can be found in Supplemental Data SD3.

### MIP capture protocol

MIP pools were prepared as described (Turner *et al.* 2009). Briefly, MIPs were pooled in equimolar amounts and treated with 50 units of polynucleotide kinase (PNK, from NEB) for 45 minutes at 37° and then 20 minutes at 80° in a 100ul reaction. The 5’- phosphorylated probes were diluted to 330nM for use in later steps. MIP libraries were based on Hiatt et al. 2013. Annealing reactions containing 500ng of target genomic DNA, 330 fmoles of MIP pool, and 1X Ampligase buffer (Epicentre) in 10ul were treated for 3 minutes at 98°, 30 minutes at 85°, 60 minutes at 60° and 120 minutes at 56°. To gap fill the product, 300 pmoles dNTPs, 7.5 umoles Betaine (Sigma), 20 nmoles NAD+, 1X Ampligase buffer (Epicentre), 5 units Ampligase, and 2 units Phusion DNA polymerase (NEB) were added to the 10ul anneal reaction and incubated for 120 minutes at 56°, and 20 minutes at 72°. To degrade genomic template and any remaining linear MIPs, 20 units Exonuclease I (NEB) and 50 units Exonuclease III (NEB) were added and incubated for 45 minutes at 37°, and 20 minutes at 80°. 10ul of this capture reaction was then amplified by 18 rounds of PCR (15 seconds at 98°, 15 seconds at 65°, 45 seconds at 72°) with 1 unit Kapa Hifi Hotstart TAQ, 10 nmoles dNTPs, and 25 pmoles each of forward and reverse primers in a 50ul reaction. Libraries were then size selected between 250-450bp and purified with Agencourt AMPure XP beads before sequencing with Illumina sequencing technology.

### MIP sequencing analysis

Sequencing data for each library was analysed using custom scripts written in R. Briefly, each sequenced library was analysed for exact matches to the expected MIP ligation arm sequences at their supposed positions within the read. This subset of matched ligation arm reads was then analysed for specific matches to the expected reference or mutant gap-fill sequence. This set was then trimmed by filtering the quality score at the specific SNV positions to include only those with Phred score >=Q30. In addition, any duplicated molecular barcodes from this set were analysed to produce a single consensus SNV read. If a duplicated molecular barcode could not find a majority agreement across gap-fill reads, it was removed from further analysis. This uniquely identified final set was used to count reference and SNV representation to produce a percentage calculated using SNV/(reference + SNV) counts. Scripts used in this analysis are available upon request.

### Normalization of MIP data

Each MIP was analysed at multiple dilutions to generate a series of curves to account for the difference between any single raw SNV-MIP call and the average of the combined SNV-MIP calls at each site. Normal distributions for each MIP were generated to fit these expected offset curves (height (h), sigma (σ), mean (μ), and offset (o)). These parameters (**Supplemental Data SD4**) were then used to re-adjust raw SNV percentages to within the expected values using the formula

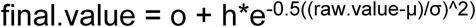

### Mapping strain determination

Based on Million Mutation Project sequencing data, a group of candidate strains was chosen to identify an appropriate mapping strain. We chose candidates with a minimum SNV density of 500±500 kbp while also minimizing for the occurrence of deleterious alleles (nonsense alleles, high Grantham scores, frameshift mutations etc.). This analysis yielded 28 candidate strains, which were then phenotyped against our reference N2 strain (VC2010) for growth, fecundity and developmental defects. The Million Mutation Project strain VC20019 exhibited growth and development similar to N2 and was chosen as a mapping strain.

### sma-9 bulk segregant mapping

*sma-9(tm572)* mutants were backcrossed twice to N2 to remove extraneous mutations before crossing with VC20019 males. Wild type phenotype F1 hermaphrodites were picked to new plates and allowed to self-cross. F2 progeny with the small body (*sma*) phenotype were then identified and pooled in groups of varying sizes to grow on 100mm NGM plates until starved. Pools were isolated for genomic DNA and subsequently analysed using MIPs.

### Temperature sensitive competitive fitness mapping

*hlh-1* and TS mutant mapping was conducted using a competitive fitness assay followed by bulk segregant analysis. Briefly, mutant animals were crossed with either VC20019 or DM7448 (VC20019;Ex[*pmyo-3::YFP*]) males. These crosses were then grown at 15° for 24-48 hours before shifting to 23° or 26°. Plates were allowed to starve and an agar chunk was transferred to fresh 100mm OP50-seeded plates. Agar chunk size was varied with 4cm^2^ pieces having the best mapping results.

Alternatively, non-starved surviving F1s were individually chosen and plated to fresh 100mm plates. When a population starved, progeny were sub-sampled by chunk-transfer to fresh media. This process continued for at least 6 rounds at the non-permissive temperature, with each transfer generally producing a new generation of animals. The remaining animals on each plate after a transfer were harvested for genomic samples used in subsequent MIP analyses.

### Temperature sensitive mutant selective bulk segregant mapping

TS mutants were crossed with DM7448 males and allowed to grow at 15°. F1 cross-progeny were chosen based on positive YFP expression. F2 animals were transferred singly to 96-well plates (liquid culture) at 15° and grown to starvation. Each well was subsampled and tested for growth at the non-permissive temperature in liquid culture. Populations that failed to thrive at the non-permissive condition were identified and pooled together from the original 15° populations. This pool was then harvested for genomic DNA used in subsequent MIPs analyses.

### Data availability

Strains are available upon request. File SD1 contains strain information including lab origin, allele name, and any available phenotype information. File SD2 contains all variant call information from the set of 173 strains described in this manuscript. Raw sequencing files are available from the NCBI Sequence Read Archive (http://www.ncbi.nlm.nih.gov/sra) under accession number SRP018046. File SD3 contains all of 89 MIP sequences and genomic targets used in the mapping analysis presented in the manuscript as well as all primer sequences required for library generation and usage on an Illumina-based sequencer. Sample MIP pools for capture can be made available upon request as well. File SD4 contains all of the required information for MIP normalization during the processing and analysis of sequence data. Custom scripts used to analyse sequencing data are also available upon request.

## Results

### Sequencing a collection of embryonic lethal TS mutant strains

We collected a total of 173 embryonic lethal TS mutant strains from three different sources that were generated either in an N2 or *lin-2(e1309)* background using different mutagenesis protocols (EMS or ENU) (Kemphues *et al.* 1988a; Ehmke *et al.* 2014) (see also Methods). After mutagenesis, candidate strains were identified through screens for embryonic lethality phenotypes at temperatures of 25°C or 26°C (**Table 1, Supplemental Data SD1**). More specifically, the 72 strains in the first set are comprised predominantly of mutant strains with early embryo defects not primarily affecting cell division or with lethality occurring after the 4-cell stage, a small number of strains with some form of sterility, and a small group of strains with low penetrance lethality or unconfirmed TS phenotypes. 67 strains make up the second set and encompass a wide range of TS phenotypes across seven broad classes from general lethality to tissue-specific developmental defects (**Table 1**). The third set consists of 34 strains identified from a mutagenesis screen for maternal effect lethality. Combined, these sets cover a diverse range of phenotypes that could reveal information on essential genes in a variety of pathways.

**Table 1.**
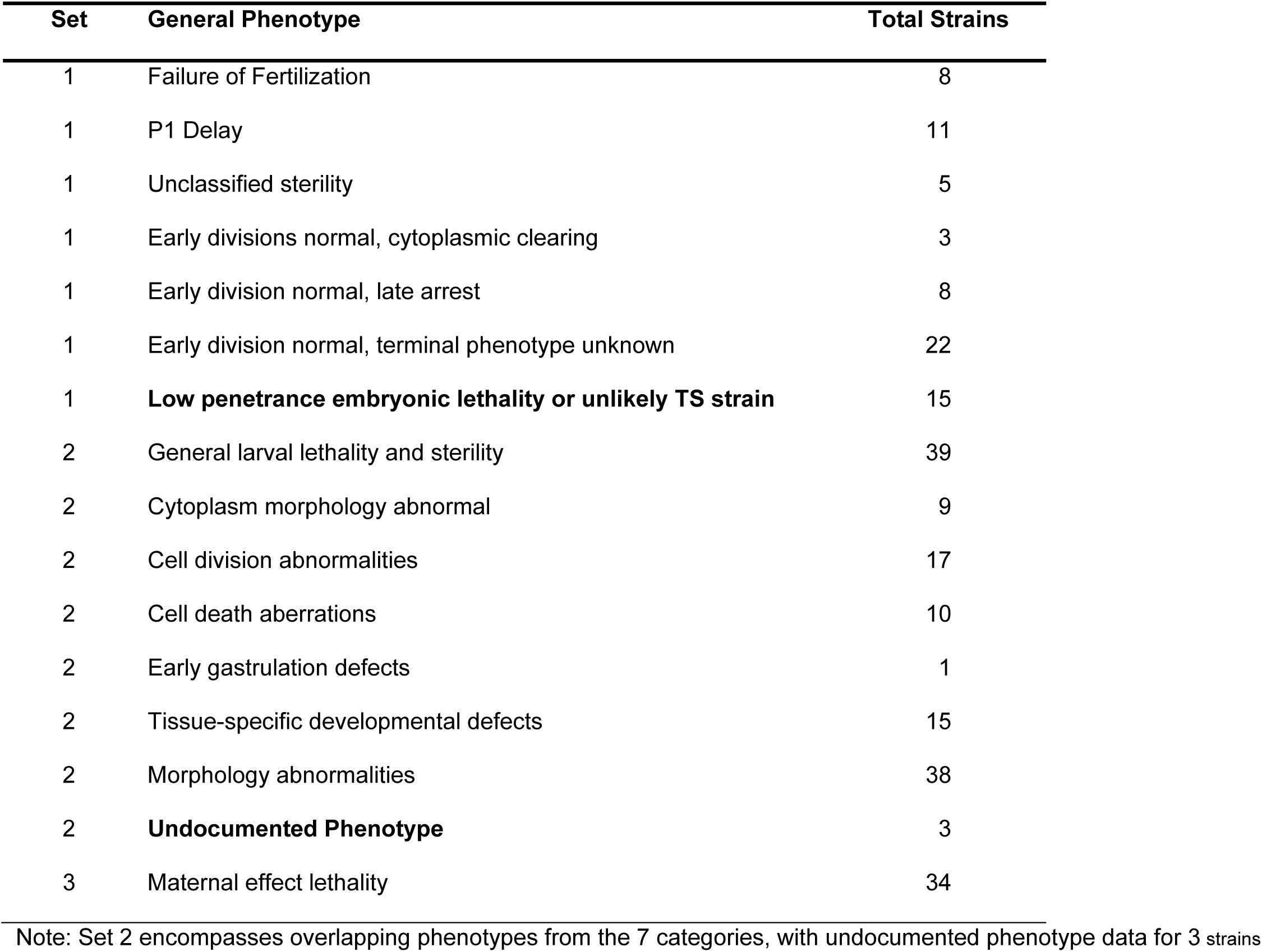
Temperature-sensitive collection phenotype summary

Prior to mapping, we sequenced genomic samples for each strain to a target minimum of 12-fold coverage (with a mean of approximately 14.5-fold coverage). Using the bioinformatic pipeline developed for the Million Mutation Project, we find that on average, each strain carries 328 SNVs including 70 missense changes, nearly 3 nonsense mutations, and approximately 1 splice site mutation per strain (**Table 2**), or < 1 change in protein coding potential per megabase of sequence within the *C. elegans* genome. Among these changes, missense mutations are the most likely to be associated with temperature-sensitive phenotypes (O’Rourke *et al.* 2011a; Lowry *et al.* 2015) although in rare instances other alterations may be responsible (Harfe *et al.* 1998). We have included a copy of the final SNV call data as part of our supplemental data (**Supplemental Data SD2**).

**Table 2.** Mutation summary from 173 strains

| Set<br>(Unique Genes) | Total<br>SNVs | Coding exons |  |  | Non-coding<br>exons (UTRs) | Introns |  | ncRNAs | Intergenic |
| --- | --- | --- | --- | --- | --- | --- | --- | --- | --- |
|  |  | Missense | Nonsense | Synonymous |  | Splicing | Other |  |  |
| Set 1<br>(4,579) | 9,285 | 1,971 | 72 | 678 | 316 | 42 | 2,770 | 131 | 3,302 |
| Set 2<br>(11,943) | 38,965 | 8,291 | 356 | 3,396 | 1,087 | 142 | 11,078 | 595 | 14,020 |
| Set 3<br>(4,269) | 8,612 | 1,896 | 78 | 723 | 251 | 35 | 2,334 | 124 | 3,171 |
| All strains | 56,803 | 12,144 | 505 | 4,793 | 938 | 217 | 16,164 | 850 | 20,478 |
| Unique Genes | 14,327 | 7,972 | 493 | 3,937 | 1,497 | 215 | 7,623 | 779 |  |

Given the challenges in *de novo* approaches to identifying the causative alleles from this collection, we turned to genetic mapping to narrow the list of candidates. We sought to develop a method that 1) produced a linkage interval at a resolution that was sufficient to limit the list to a handful of candidates and was comparable in resolution to current methods; 2) replaced the Hawaiian isolate as a mapping strain with one more genomically similar to N2; 3) allowed for a cost-efficient high-throughput workflow. We postulated that mapping with smMIPs in a targeted sequence capture strategy (Hiatt et al., 2013) would provide similar mapping resolution to current WGS methods, allow the use of a strain minimally divergent from N2 and decrease the overall sequencing burden per sample.

### Targeted sequencing of select polymorphisms by MIP-MAP

We re-engineered the original probe structure to adapt the single molecule targeting capabilities of smMIPs for analysing SNV abundance as a method of mapping (hereby referred to as MIP-MAP) (**Figure 1**). Each MIP is composed of an 80bp oligonucleotide with a pair of annealing arms at the 3’ (extension arm) and 5’ (ligation arm) ends totalling 40 basepairs. Between these two arms is a molecular barcode sequence – randomized for each individual oligonucleotide – as well as a common MIP backbone used in library preparation and amplification steps (**Figure 1A**). Each MIP is designed to target a locus where a known variant is located within the first 18 basepairs of the 100- 150 basepairs of gap-fill sequence (**Figure 1AB**). The gap-fill sequence also serves to unambiguously map the recovered sequence. The library is then amplified, at which time an experimental index sequence is incorporated (**Figure 1C**). This design places all the vital information within a single-end 50 basepair Illumina sequencing read (**Figure 1D**). After sequencing, each sample library samples is demultiplexed based on an experimental index after which molecular barcodes and gap-fill sequence data are used to eliminate PCR duplicates (**Figure 1E, Methods**). The remaining unique sequencing information is used to determine a MIP target’s SNV representation versus the total reads matching that locus as calculated by

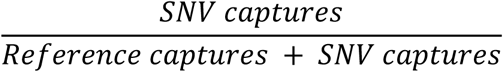

**Figure 1.**
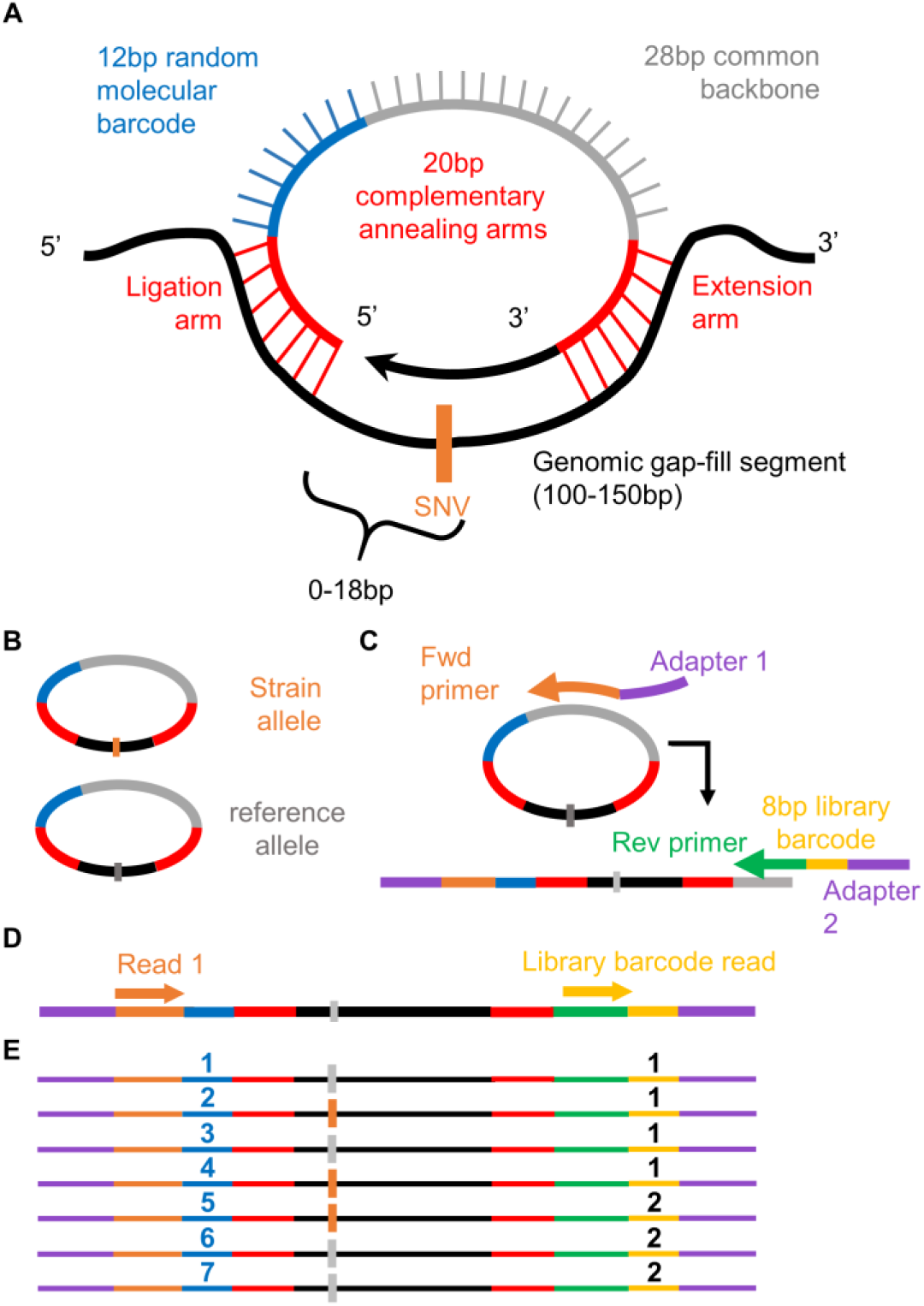
MIPs design and workflow. The design of the single-molecule Molecular Inversion Probes (smMIPs, Hiatt et al., 2013) was modified to relocate the molecular barcode. Overall the MIP design incorporates two 20bp annealing arms, a 12bp molecular tag and 28bp common backbone. After annealing to a target segment ranging in size from 100-150bp (A), the MIP is gap-filled with high-fidelity polymerase capturing the sequences of interest and circularized via ligation (B). The SNV of interest is located within an 18bp gap-fill window upstream of the ligation arm. Uncircularized DNA is then degraded via exonucleases. The remaining MIPs are linearized by combining with sequencing adapters and library barcodes during PCR amplification (C). A single end read captures both the molecular tag, 5’ annealing sequence, and enough genomic sequence to confirm correct target capture. This information is then demultiplexed (E) and used as a means to compare sequence variant ratios.

With a targeted sequencing strategy in hand, we next examined the parameters that would influence our selection of probes across the *C. elegans* genome: mapping resolution, mapping strain choice, and MIP accuracy. Mapping resolution is generally limited by the number of F2 animals pooled rather than SNV density. With just 100 F2s picked, average resolution is no more than 1 cM or about 1-3 Mb depending on the region of the genome (Barnes *et al.* 1995; Rockman and Kruglyak 2009; Doitsidou *et al.* 2010). Accordingly, sampling ~100 SNV markers across the genome would match the expected genetic resolution. At this density, two hundred thousand unique sequencing reads would provide deep coverage of each SNV, allowing the accurate estimation of even relatively rare linked SNVs – a vast improvement in sequencing burden versus current paradigms.

Decreasing the density requirement of our mapping SNVs also permitted us to survey the Million Mutation Project strains for potential alternatives to the Hawaiian strain. The Million Mutation Project strains average ~400 mutations each, yet many are roughly wild type in appearance. We screened for a collection of strains that carried minimal numbers of predicted deleterious or missense coding variants and a relatively even distribution of SNVs across the genome at a frequency of at least 1 per 500 kilobases. From 28 candidates, we identified a strain, VC20019, with developmental timing and fecundity similar to that of our N2 wild type strain (VC2010). VC20019, also referred to as the mapping strain, has a total of 269 mutations, 3 nonsense, and 51 missense alleles, thus having enough genomic diversity compared to N2 to meet our molecular marker density requirements while having a reduced likelihood of potentially negative interactions.

We designed MIPs targeting 96 SNV sites across the VC20019 genome, generating intervals spaced an average of 0.98 ± 0.36 Mbp apart (**see Supplemental Data SD3**). To identify the efficiency of annealing and overall fidelity of these probes, we generated a series of VC20019 and N2 genomic DNA mixtures for capture reaction and sequencing. From these libraries, we identified sequence-specific biases for each MIP and generated normalization curves for subsequent capture/sequencing reactions (**Supplemental Figure S1, Supplemental Data SD4, Methods)**. We removed seven VC20019 MIPs that consistently produced low sequencing read counts, high ratios of non-specificity, or unpredictable allelic biases, bringing our total probe count to 89 probes (1.06±0.43 Mbp intervals).

We also considered that the frequency for linked and therefore rare SNVs could be adversely impacted by false-positive MIP-MAP reads. To gauge the impact of false-positives on the accuracy of our calls, we designed an additional 176 MIP-MAP targets based on SNVs in 44 strains from the Million Mutation Project collection. We analysed the final SNV representations of each MIP capture library from a genomic DNA mixture that excluded the selected Million Mutation Project target strains. We sequenced this set of MIPs to an average depth of ~35K reads. We removed data from poorly amplifying MIPs, leaving 174 MIP target loci that were expected to have 0% mutant SNV representation. MIP reads with positive mutant calls must originate from gap-fill, PCR, or sequencing errors and the overall percent mutant SNV representation was calculated as the false-positive rate. As a further quality analysis step, we filtered out low-quality sequencing reads at specific SNV sites and calculated the average false positive rate was 0.0122% ± 0.0129%. (**Supplemental Figure S2A, Methods**).

Lastly, we determined the sequencing efficiency of the MIP-MAP method by comparing the population of unique molecular barcodes for each MIP versus the total reads identified for that MIP. For each MIP in our false positive set of 174 MIP targets, the mean number of sequencing reads per unique molecular barcode was 1.05 with a mean depth of 35,000 reads per MIP. We analysed an additional 22 libraries from another sequencing run, consisting of the 89 mapping strain MIPs in each library. The mean reads per barcode across 1,958 data points was 1.01 with a mean depth of 7000 reads per MIP (**Supplemental Figure S2B**). These results suggest that we are capturing a nearly 1:1 representation of the genomic sample, with few redundant reads from non-unique capture sequences.

In summary, we re-engineered the smMIP format to produce MIP-MAP which targets the genome with high fidelity and high sequencing efficiency at a mapping resolution comparable to current methods. We identified a polymorphic strain similar to N2 with no observed adverse phenotypes and a reduced likelihood of negative genetic interactions. We next turned to examining the MIP-MAP protocol in mapping known mutant alleles and then uncharacterized strains of varying difficulty, such as those from the TS collection.

### Mapping sma-9 and hlh-1 mutations using MIP-MAP

To investigate the mapping capabilities of the MIP-MAP method, we first focused on mapping with two distinct phenotypes; a clearly identifiable *sma-9* small body mutant and the well-characterized *hlh-1* TS lethal mutant. We proceeded by using the LW478, *sma-9(tm572)*X, strain in a standard bulk segregant strategy similar to WGS (**Figure 2A**). Briefly, we crossed *sma-9(tm752)X* hermaphrodites with mapping strain males and from F1 cross progeny chose groups of F2 *sma* phenotype animals (10, 25, 50, 75, 100, 200) to grow until starvation on OP50-seeded 100mm NGM plates before isolating genomic DNA for MIP-MAP capture reaction and sequencing (**Figure 2, Supplemental Figure S3**). Our expectation with such a strategy was to observe a region where VC20019 SNVs were nearly depleted due to linkage disequilibrium in selecting for the *sma-9* allele from the mutant strain. We generated MIP-MAP data by growing populations seeded with 10 Sma phenotype F2s. These replicate samples had more background signal across the genome than subsequent experiments and our interval of interest was 1.5Mbp and 4.2Mbp in these samples (**Figure 2B**). Seeding populations with 75 Sma phenotype F2s yielded similar mapping profiles that narrowed the likely interval to 2.6Mbp (**Supplemental Figure S3C**). Likewise, two mapping sets seeded with 200 F2 animals also consistently identified a 2.6Mbp interval containing *sma-9* (**Figure 2C**). Upon closer inspection of our *sma-9* mapping intervals, we observed two sites ~590kb apart that typically share nearly identical SNV representation, suggesting linkage disequilibrium. It is not surprising, therefore, to find that *sma-9* is located equidistantly between these points, making it less likely to generate a sharp single-point peak even with a high number of F2s as input (**Figure 2D**). On average, the MIP targets in our 75-F2 sets had 3778 unique reads, suggesting that each F2 haplotype was sampled an average of 25 times. Therefore, the MIP-MAP technique can sufficiently capture an accurate representation of the recombinant landscape across the population.

**Figure 2.**
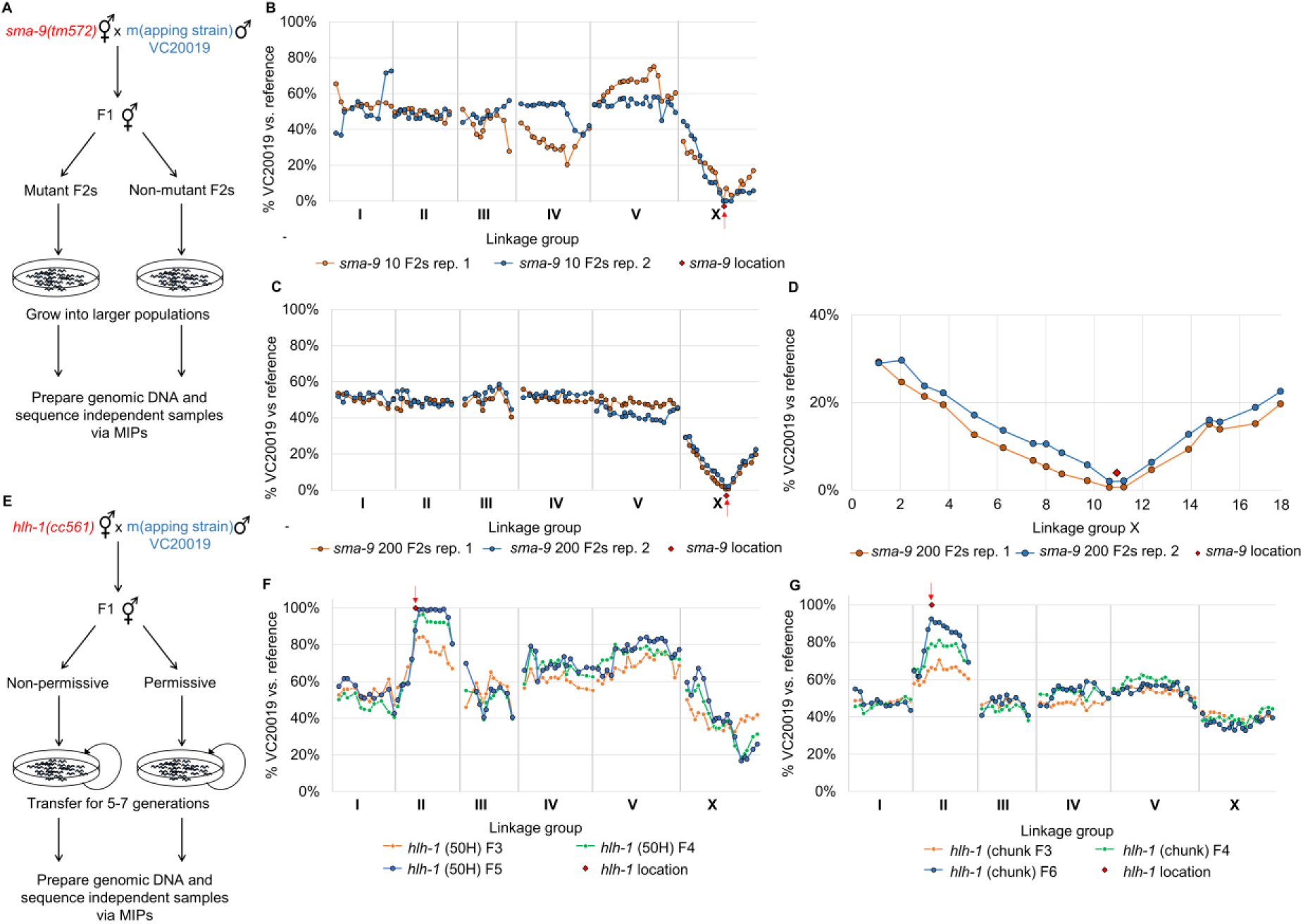
Mapping of *sma-9* and *hlh-1* via VC20019 and MIP-MAP sequencing. Bulk segregant mapping scheme for *sma-9* (A) using MIPs that target a number of VC20019- specific SNVs across the genome. Two replicate MIP-MAP samples of populations grown from 10 sma phenotype F2 animals sufficiently identified a *sma-9*-associated interval on X within a 1.5Mbp to 4.2Mbp window (B) and replicates using 200 sma F2 recombinants produced a 2.6Mbp window (C). Closer examination of the MIP target sites on LGX show *sma-9* is located equidistantly between two probes only 590kb apart (D). The *hlh-1(cc561)* temperature sensitive embryonic lethal allele was mapped using a competitive fitness mapping method (E). PD4605 hermaphrodites were mated with VC20019 males and non-permissive temperatures were used to select against the subsequent *hlh-1(cc561)* homozygous progeny. Starting with F1 cross progeny and transferring mixed-stage sub-populations of animals (F,G) for up to 7 generations, a region corresponding to the *hlh-1* locus on LGII was successfully identified by the fixation of VC20019-specific SNVs. A mapping interval of 2.8-7.8Mbp was identified by the F5 generation when transferring populations of 50 L1s (F). A 2.7Mb interval interval was observed by the F6 generation when transferring 4cm^2^ chunks of mixed-stage animals (G). Each line present in (F) and (G) represents mapping data from a different generation for the same biological replicate.

We next explored if MIP-MAP was applicable to more challenging mutant phenotypes. For instance, isolating homozygotes of non-conditional lethal/sterility alleles or identifying the correct homozygous F2 populations of low penetrance alleles may not always be feasible. We approached this scenario with a competitive fitness mapping assay (**Figure 2E**). We hypothesized that we could exploit the fitness defect of an allele during the mapping process and gradually replace it with a more “fit” version from the mapping strain. Using the *hlh-1(cc561)*II temperature sensitive embryonic lethal allele, we attempted to passively map its locus by using non-permissive temperatures to select against this allele over several generations. In contrast to the *sma-9* mapping, we expected to identify a region of interest by the fixation of VC20019 SNVs towards 100% representation within the mapping profile. We refined the parameters of the competitive fitness MIP-MAP method by varying the number of progeny passaged at each generation. Briefly, we crossed *hlh-1* animals to VC20019 males at 15°C on 50mm NGM plates for approximately 24 hours before shifting these plates to between 23°C and 26°C for the remainder of the experiment. We expected only cross progeny F1s would thrive at this temperature. F1s were picked and self-fertilized before randomly picking L1- or L2-staged F2 larvae. With each subsequent generation, a subpopulation (50, 100, 200 or 400 animals) was passaged to a new plate for expansion (**Figure 2F, Supplemental Figure S4A-C**). We continued in this manner for 4-7 generations, collecting samples for MIP-MAP library preparation and sequencing at each passaging.

Based on our observations from mapping *sma-9*, we hypothesized that at least 75 F2 animals (upwards of 150 recombinant chromosomes) homozygous for our region of interest, were required to consistently generate the smallest mapping interval compatible with the density of our markers. The mapping profile from a competitive fitness assay might therefore be influenced by the number of unique recombinant chromosomes that carry our region of interest in each generation. We reasoned that subsampling in small numbers could limit recombinant representation and increase the incidence of bottlenecking within the population. A chance reduction in recombinant chromosome diversity could lead to quick fixation of a region of interest but with reduced resolution. Conversely, subsampling in larger numbers could give a greater range of recombinant genomes, reduce bottlenecking and generate better resolution while likely requiring more time to reach fixation of the more “fit” wild type region.

With this in mind we varied our passaging size and observed that an overrepresented region of VC20019-associated SNVs on LGII with varying degrees of resolution (**Figure 2FG**). This candidate interval included the location of *hlh-1* and suggested that we had correctly mapped the *cc561* allele. The smallest possible mapping intervals encompassing the location of *cc561* are 1.7 and 1.9Mbp. When passaging smaller populations (i.e., 50 animals per generation), fixation occurred quickly but in some cases resulted in a larger mapping interval across LGII (**Figure 2F, Supplemental Figure S4A**). The resolution of the MIP-MAP approach, however, improved in the populations seeded with ≥ 200 randomly chosen F2s (**Supplemental Figure S4BC**). Thus, sampling more animals generated the optimal minimal mapping interval of 1.71 Mbp with only a one to two generation increase required for fixation. To exclude the possibility of a sampling bias for healthier animals when hand-picking our *hlh-1* mapping recombinants, we replicated these experiments with an alternate procedure by transferring completely randomized populations via agar chunks of starved animals for 5-9 generations (**Figure 2G**, **Supplemental Figure S4D-F**, **Methods**). We again observed that larger transfer sizes associated with better resolution but this approach also generated a more pronounced delay of 2-8 additional generations before fixation.

We further validated the competitive fitness assay by investigating the general fitness of the mapping strain in a cross to the reference strain VC2010 and growing cross-progeny at a range of temperatures. We observed only partial fixation of LGV at high temperature over 8 generations but other loci preferences within the genome appeared negligible (**Supplemental Figure S5A-C**), thereby confirming that our *hlh-1* results were not an artefact of the mapping strain used. We could therefore reliably identify a strong temperature-sensitive lethal allele through a simplified competitive fitness selection process. In addition, we noted the weak beginnings of fixation on LGII when growing an *hlh-1* mapping population at 15°C (**Supplemental Figure S6**), suggesting a small population growth defect was conferred by the *cc561* allele at the so-called permissive growth temperature. We next turned to see if the MIP-MAP competitive fitness assay could be applied to genetically mapping uncharacterized TS lethal alleles.

### Using MIP-MAP to identify TS mutations that confer lethal phenotypes

Our success with *hlh-1* led us to hypothesize that we could, with good resolution, map mutant alleles from our large collection of sequenced TS lethal strains. Unlike our previous trials, these mutants carry hundreds of variants – any one of which may influence the outcomes of a competitive fitness assay. Furthermore, TS alleles of low or weak penetrance could also prove troublesome given a reduced selection coefficient during the propagation process. Regardless of these factors, from the collection of TS mutants we chose 15 strains (**Table 3**) of high sequencing depth but varying phenotypes to attempt genetically mapping with MIP-MAP.

**Table 3.** Table of temperature-sensitive mutants mapped

| Strain | Set | WGS Coverage | Phenotype Penetrance | Timing of lethality and phenotype observations at 26°C |
| --- | --- | --- | --- | --- |
| VC50022 | 3 | 16.6 | 100% | Arrest in early embryogenesis |
| VC50028 | 3 | 40.7 | 100% | Sterility of P<sub>0</sub> when plated at L4. Any F1s produced arrest at early larval stages. Some sterility also noted at permissive temperatures. |
| VC50031 | 3 | 16.4 | 100% | Sterility of P<sub>0</sub> when plated at L4. Any F1s produced arrest at early embryogenesis. |
| VC50141 | 3 | 16.9 | 100% | Sterility of P<sub>0</sub> when plated at L4. Any F1s produced arrest at early embryogenesis. |
| VC50174 | 1 | 21.9 | <80% | Sterility of P<sub>0</sub> when plated at L4, but also have bag phenotype so F1 are still produced. Failure of fertilization with vacuolated spermatozoa observed. |
| VC50178 | 1 | 20.7 | 100% | P<sub>0</sub> when plated at L4 still produce progeny but F1 population is observed to be sterile. Severe morphogenesis defect, possible 2-fold arrest. |
| VC50182 | 1 | 19.8 | 100% | Early divisions normal, late arrest. Mostly 2-fold arrest with movement. |
| VC50255 | 1 | 20.0 | 100% | Sterility of P<sub>0</sub> when plated at L4 with bag phenotype of dead eggs likely due to embryonic arrest. Any F1 produced are sterile. P1 delay is observed in embryos. |
| VC50260 | 1 | 22.9 | 99% | Sterility of P<sub>0</sub> when plated at L4 with bag phenotype of dead eggs likely due to embryonic arrest. Any F1 produced are sterile. P1 delay is observed in embryos. |
| VC50352 | 2 | 23.4 | <100% | Slightly leaky as P<sub>0</sub> can give rise to F1, and then F2 but these are usually sickly and sterile. In some cases they can make F3. Pretzel inviable, asynchronous cell cycle, later stage uncoordinated. |
| VC50360 | 2 | 22.2 | 100% | P<sub>0</sub> have a mild roller phenotype, sluggish in appearance. F1 progeny appear to arrest in early embryogenesis. Pretzel inviable, pretzel deformed, cell adhesion/migration defect. |
| VC50374 | 2 | 20.9 | 99% | P<sub>0</sub> when plated at L4 produce unhatched eggs of various forms and appears embryonic lethal.<br>Any F1s produced are sterile. Arrest of development at premorphogenetic stage |
| VC50375 | 1 | 19.0 | 100% | F1 sterility with P1 delay observed. |
| VC50380 | 2 | 19.6 | 100% | P<sub>0</sub> when plated at L4 appear active and produce eggs. F1 progeny arrest in late embryogenesis and L1. Pretzel inviable, abnormal cytoplasm morphology, spindle defects, aberrant pharyngeal cluster and pretzel deformed. |
| VC50383 | 2 | 21.7 | 100% | P<sub>0</sub> plated at L4 appear active but produce eggs that fail to hatch. Pretzel inviable, larval lethal, and muscle defects |

Since these highly mutated strains could harbor mutations that reduce brood size or mating efficiency, we modified our process to ensure the picking of cross-progeny by crossing our TS mutants with DM7448 (a VC20019 strain carrying an extrachromosomal *pmyo-3*::YFP marker) males and picking YFP-positive F1 hermaphrodites. These F1s were subjected to the previously-described competitive fitness assay via chunk- or wash-transfer and in some instances we supplemented our analysis with a multi-well liquid format of bulk segregant mapping for clarification (**Figure 3A-D, Methods**). Our mapping results (**summarized in Figure 3E**, **Table 4**) can be categorized into increasing levels of analysis difficulty: clear single TS loci (**Figure 4AB, Supplemental Figure S7**); single TS loci with additional partial fitness defective loci (**Figure 4CD, Supplemental Figure S8**); and multi-locus profiles resolved by additional investigations (**Figure 4E-G, Figure 5A-C, Supplemental Figure S9**).

**Figure 3.**
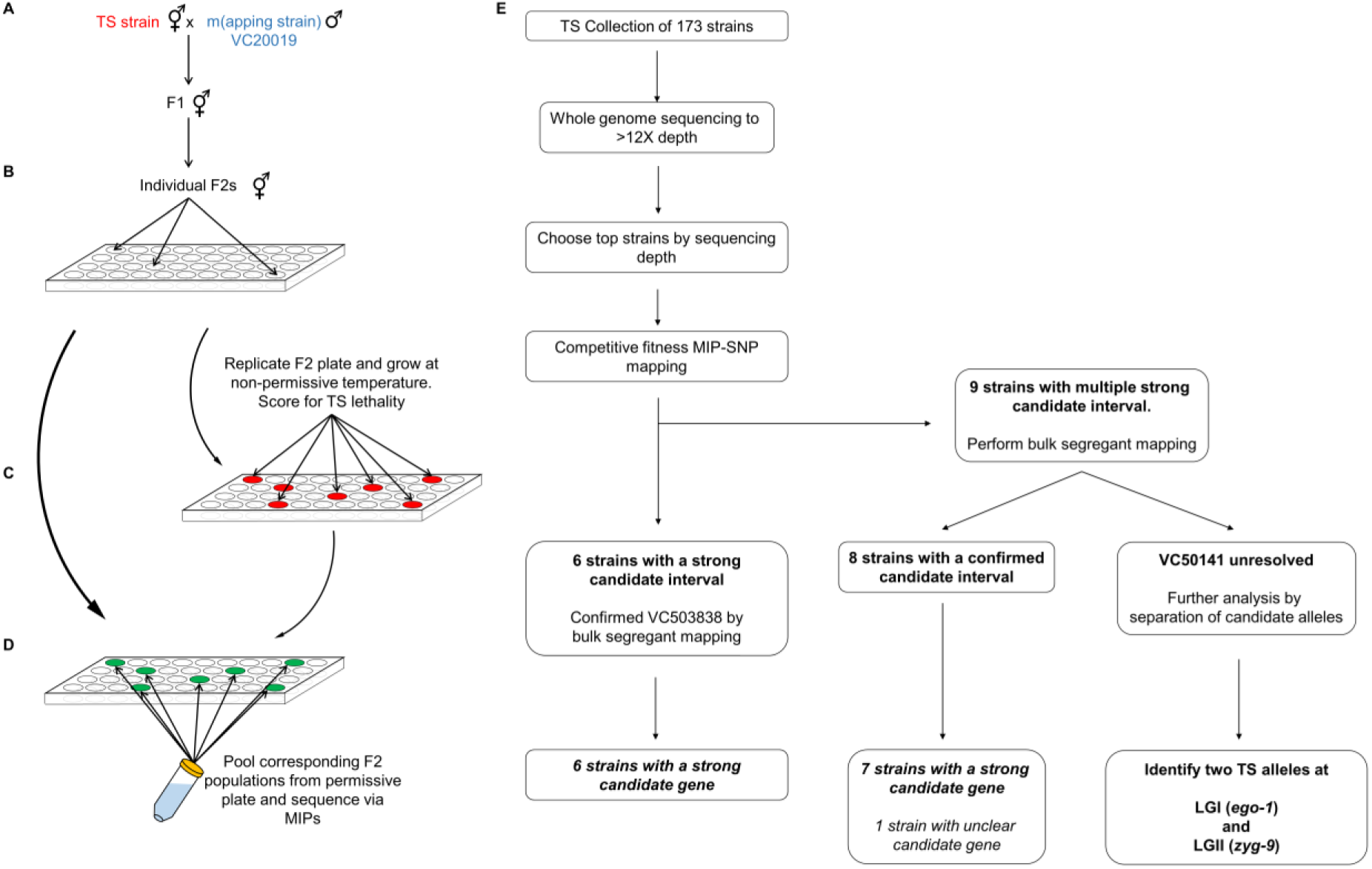
Workflows for liquid bulk segregant mapping and TS mutant analysis. (A) VC20019 males were mated to a TS mutant strain with cross progeny F1 animals used to generate F2 recombinants. Single F2s were used to seed wells (B), growing these populations at permissive temperatures before replicating them to grow in non-permissive conditions (C) and identify populations that failed to thrive. Dead populations (red) were then chosen from the original wells (green) and pooled together (D) to prepare a single genomic sample for MIP-MAP sequencing. Samples from multiple plates were combined in the mapping of a single mutant depending on allele penetrance or the number of positive F2 wells identified. The workflow (E) of mapping TS mutants shows 6 strains could be mapped by competitive fitness MIP-MAP alone; 8 strains benefitted from additional mapping via the liquid bulk segregant protocol; and 1 strain (VC50141) required additional analyses.

**Figure 4.**
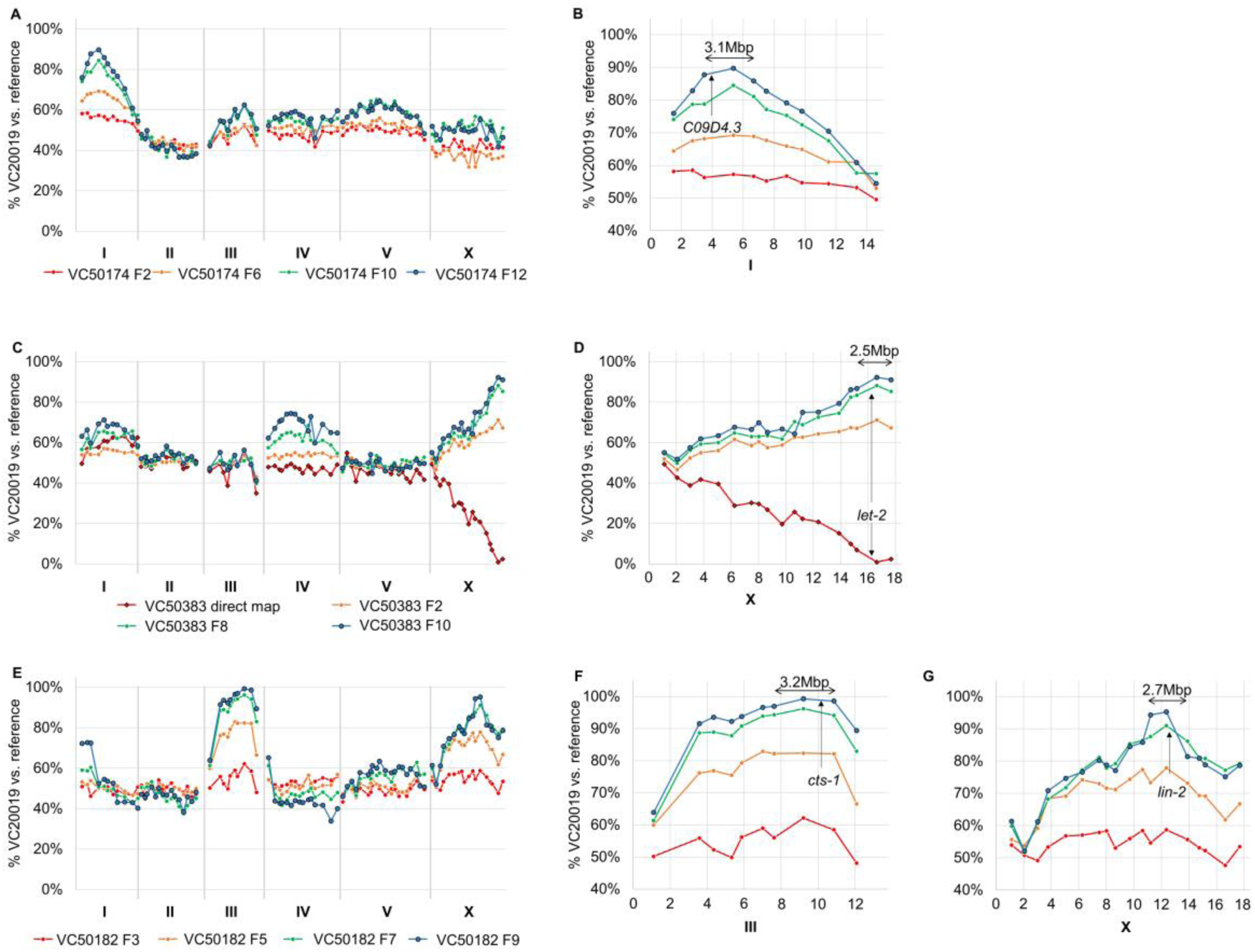
Successful mapping of temperature-sensitive lethal mutant strains via competitive fitness and liquid-format MIP-MAP techniques. 16 TS lethal mutants with genomic sequencing data were mapped by the competitive fitness approach used with *hlh-1*. Mapping results yielded a diverse group of mapping profiles including clean single-locus mappings as seen in VC50174 (AB); profiles such as VC50383 with small additional loci or peaks appearing at later generations but resolved by additional liquid-format bulk segregant mapping of phenotypes (CD); and multiple-locus profiles resolved by identifying shared background mutations in related strains as in VC50182 (EF). Each line present in a graph represents mapping data from a different generation for the same experimental replicate or the results of a liquid-format bulk segregant mapping experiment (indicated as direct map with diamond points).

**Figure 5.**
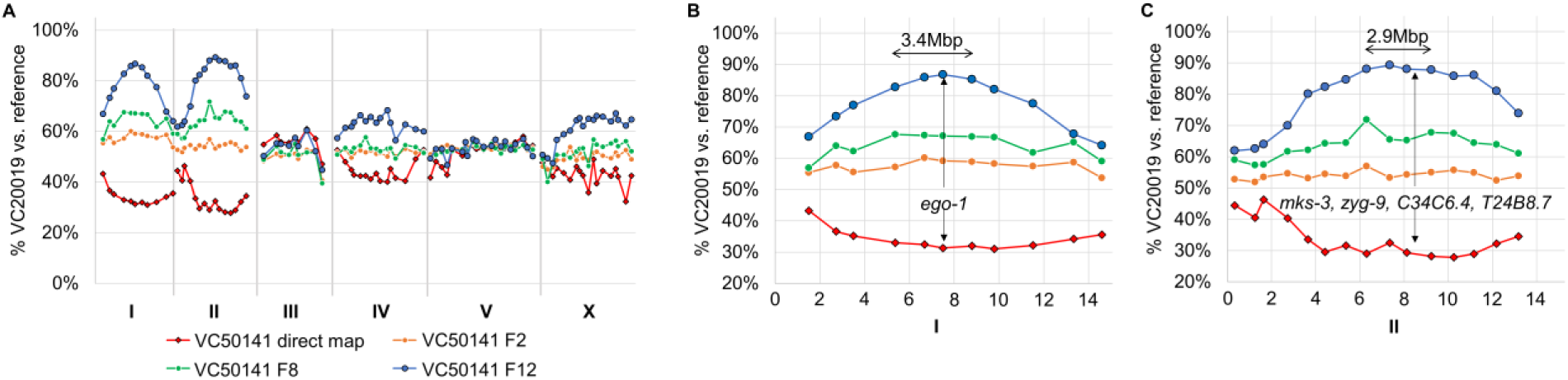
The TS strain VC50141 has two independent TS alleles of *ego-1* and *zyg-9*. Both competitive fitness mapping and liquid-format bulk segregant mapping confirmed that the TS lethality in VC50141 was related to two loci (A). The first interval located on LGI contains only a single relevant coding mutation in *ego-1* (B). The second locus, appearing to be of equal penetrance on LGII contains 4 candidates, of which *zyg-9* is the most likely causative allele (C). Each line present in the graph represents mapping data from a different generation for the same experimental replicate or the results of a liquid-format bulk segregant mapping experiment (indicated as direct map with diamond points).

Three strains yielded clean, single-locus profiles with only one or two TS candidate alleles in each interval. Among these strains, VC50174 mapped to an adenine to thymine transversion resulting in an E196D mutation in the fifth exon of *C09D4.3* (**Figure 4AB**), a gene with no prior phenotype data to suggest an essential role in development. Available RNAseq data reports *C09D4.3* transcript abundance peaking at the L4 stage, most especially in males (Hutter *et al.* 2009; Boeck *et al.* 2016) and is suggested to have preferential expression in the germline (Grün *et al.* 2014). Furthermore, we observed that VC50174 animals exhibited a failure of fertilization and vacuolated spermatozoa. VC50022 mapped to an interval with a coding mutation in *tbb-2*; one of two nearly functionally redundant *C. elegans* β-tubulin genes. Although not entirely essential, multiple labs have identified semi-dominant TS-lethal alleles for this gene (Wright AJ 2003; Ellis *et al.* 2004). In particular, the *tbb-2* mutation in VC50174 is located within the same exon as a semi-dominant embryonic lethal allele *t1623* (Gönczy et al. 1999). The other possible candidate, *F56D2.5*, is at the edge of this interval, shows low expression throughout embryogenesis (Boeck *et al.* 2016) and has no useful RNAi phenotype data. VC50260 maps to an interval encompassing a single coding mutation for *npp-8* which has also been reported as an essential gene with a documented TS allele, *ax146* (Asencio *et al.* 2012).

In three instances, our TS mapping profiles suggested the presence of additional weak low-fitness alleles at secondary loci (**Figure 4CD, Supplemental Figure S8**). In these cases, we combined mapping data from multiple time points with the expectation that the TS-associated locus was the least competitive at non-permissive temperatures and would therefore fixate at a faster rate. When we were unable to ambiguously resolve the correct chromosome, we further mapped the strain with a liquid bulk-segregant assay (**Figure 3A**). By growing singled F2 populations in 96-well liquid format we could subsample each and phenotype for TS lethality, going back to the original populations later to pool for MIP analysis. This method would generally produce a low number of TS F2s (**Supplemental Table S1**) but could yield a single mapping signal across the *C. elegans* genome. Of particular interest, the strain VC50383 yielded a number of a candidates within the mapping interval. The most likely candidate, however, was a *let-2* exon 16 guanine to adenine transition resulting in a G1385E change which was confirmed through complementation testing. Coincidentally, another set 3 strain, VC50380, also mapped to an interval that included another *let-2* guanine to adenine transition (G1110E) in exon 14. *In toto*, there are 9 separate *let-2* mutations from our collection that could further elucidate the complex nature of this gene (Meneely and Herman 1981).

Our final group of nine TS strains represent multi-locus mapping profiles that fixed to similar proportions within a short period of time (**Figure 4EFG, Supplemental Figure S9**). These strains may carry additional mutations that confer disadvantages during the competitive mapping process. Again, a combination of comparing fixation rates at each locus and the liquid segregant mapping assay were used to resolve the TS-associated locus. In three of these strains, our mappings included a clearly marked interval on chromosome X (**Figure 4EFG**) which encompasses the *lin-2* gene. Strains from both set 1 and set 3 harbor the lin-2(*e1309)* mutation which causes egg retention and reduced brood size – negative influences in a competitive fitness assay. From these strains in particular, VC50182 yielded two candidates within the interval on LGIII of which *cts-1* (a citrate synthase) is the likely prime candidate. VC50182 animals exhibited late arrest at the 2-fold stage, while reported *cts-1* RNAi phenotypes include embryonic lethality and sterility (Kamath *et al.* 2003; Melo and Ruvkun 2012).

Also of note, in mapping the strain VC50141, we observed the concomitant but incomplete fixation of two loci – one each on LGI and LGII – when using our competitive fitness assay (**Figure 5A**). Similarly, the liquid-format bulk segregant assay identified the same two linkage groups at approximately 30% mapping allele representation versus an expected 0% generated from 75 TS-positive wells in 387 randomly chosen F2s. This phenomenon suggested the presence of two TS alleles. The LGI interval contained only a single coding change which corresponded to a guanine to adenine transition (E994K) in the RNA polymerase homolog *ego-1* (Qiao *et al.* 1995; Smardon *et al.* 2000) (**Figure 5B**). There are no previously reported conditional-lethal alleles for *ego-1*. The LGII locus contained 4 possible candidates including a cytosine to thymine (P632S) *zyg-9* mutation (**Figure 5C**). There are a large number of reported TS alleles located across *zyg-9* (Bellanger *et al.* 2007), suggesting it is a good candidate for VC50141. To confirm whether these two loci were independent TS alleles or if they perhaps interacted to produce a synthetic TS lethal effect, we proceeded to isolate these two alleles into separate strains (**Methods**) followed by scoring for lethal phenotypes at 26°C. Based on our observations we concluded the presence of two TS-lethal phenotypes, sterility and embryonic lethality, linked to the *ego-1* and *zyg-9* alleles respectively. Thus our mapping methods were able to identify both *ego-1(ax458)*I and a *zyg-9(ax3101)*II as the primary candidates for two independent TS-lethal mutations in this strain.

Overall, for our set of TS strains, this method yielded relatively small mapping intervals that contained greatly reduced numbers of candidate alleles (**Table 4**) - simplifying the process of finding the causative allele. Of 15 candidate strains, 6 were mapped with only the competitive fitness approach while 9 had additional bulk segregant mapping performed, with the special case of VC50141 requiring additional intervention (**Figure 3E**). In three strains, our mapping intervals contained only a single relevant coding sequence candidate, while in twelve intervals we narrowed candidates to a primary variant based on a combination of gene description, expression pattern, RNAseq, and RNAi phenotypes data from Wormbase and strain phenotype observations (**Table 5**). Of the sixteen mapping intervals identified only one (VC50352; LGIV) could not be narrowed to a primary candidate based solely on a comparison of the strain phenotype, mapping data and available literature. Of the fifteen TS intervals with primary candidate genes, six have no reported TS alleles listed in Wormbase and one (*C09D4.3*) has no previously reported embryonic lethal phenotypes. Of the other candidates with no available TS alleles, *ego-1*, *let-805*, *mog-4*, and *par-5* alleles with known phenotypes are only available from the Caenorhabditis Genetics Center as viable strains with genetic balancers. The gene *cts-1* only has a single deletion strain available which is also listed as sterile/lethal.

**Table 4.** Summary of temperature-sensitive mutant candidate mapping intervals

| Strain | Set | Mapping methods | Mapping interval | Candidate gene(s) |
| --- | --- | --- | --- | --- |
| VC50022 | 3 | CF | LGIII:3579637-5328496 | *tbb-2, F56D2.5* |
| VC50174 | 1 | CF | LGI:3494925-6676830 | *C09D4.3* |
| VC50178 | 1 | CF | LGIII:0-3579637 | *M01G5.1, let-805* |
| VC50182 | 1 | CF | LGIII:7617321-10853887 | *cts-1, enu-3* |
| VC50260 | 1 | CF | LGIV:1125542-3797513 | *npp-8* |
| VC50380 | 2 | CF | LGX:15150772-17676467 | *Iron-3, csb-1, let-2, K09E3.5* |
| VC50028 | 3 | CF and 96-well | LGII:121799849-15279345 | *cox-10, jmjd-2, K10H10.4, mog-4* |
| VC50031 | 3 | CF and 96-well | LGV:18400066-20183925 | *Y69H2.3, asp-18, emb-4* |
| VC50255 | 1 | CF and 96-well | LGIII:7617321-10853887 | *C02F5.10, pri-1* |
| VC50352 | 2 | CF and 96-well | LGIV:5057999-7478926 | *vit-2, pqn-62, Y34B4A.6, ifta-1, C09B8.8, pak-1, col-171* |
| VC50360 | 2 | CF and 96-well | LGV:11035658-14223805 | *pak-2, F35B12.9, rad-50, sqt-3, F28C1.1, egl-10* |
| VC50374 | 2 | CF and 96-well | LGIV:1125542-3797513 | *F42A6.6, hrp-1, Y37E11AL.6, map-1, taf-6.2* |
| VC50375 | 1 | CF and 96-well | LGIV:9147769-12028068 | *dct-15, K08F4.5, par-5* |
| VC50383 | 2 | CF and 96-well | LGX:15180772-17676467 | *rab-14, let-2, nas-39, acs-9, F52G3.1, gcy-11, crb-1* |
| VC50141 | 3 | CF, 96-well, | LGI:6676830-8797510 | *ego-1* |
|  |  | segregation testing | LGII:6305987-9232191 | *mks-3, zyg-9, C34C6.4, T24B8.7* |

CF = competitive fitness mapping; 96-well = liquid bulk segregant mapping

**Table 5.**
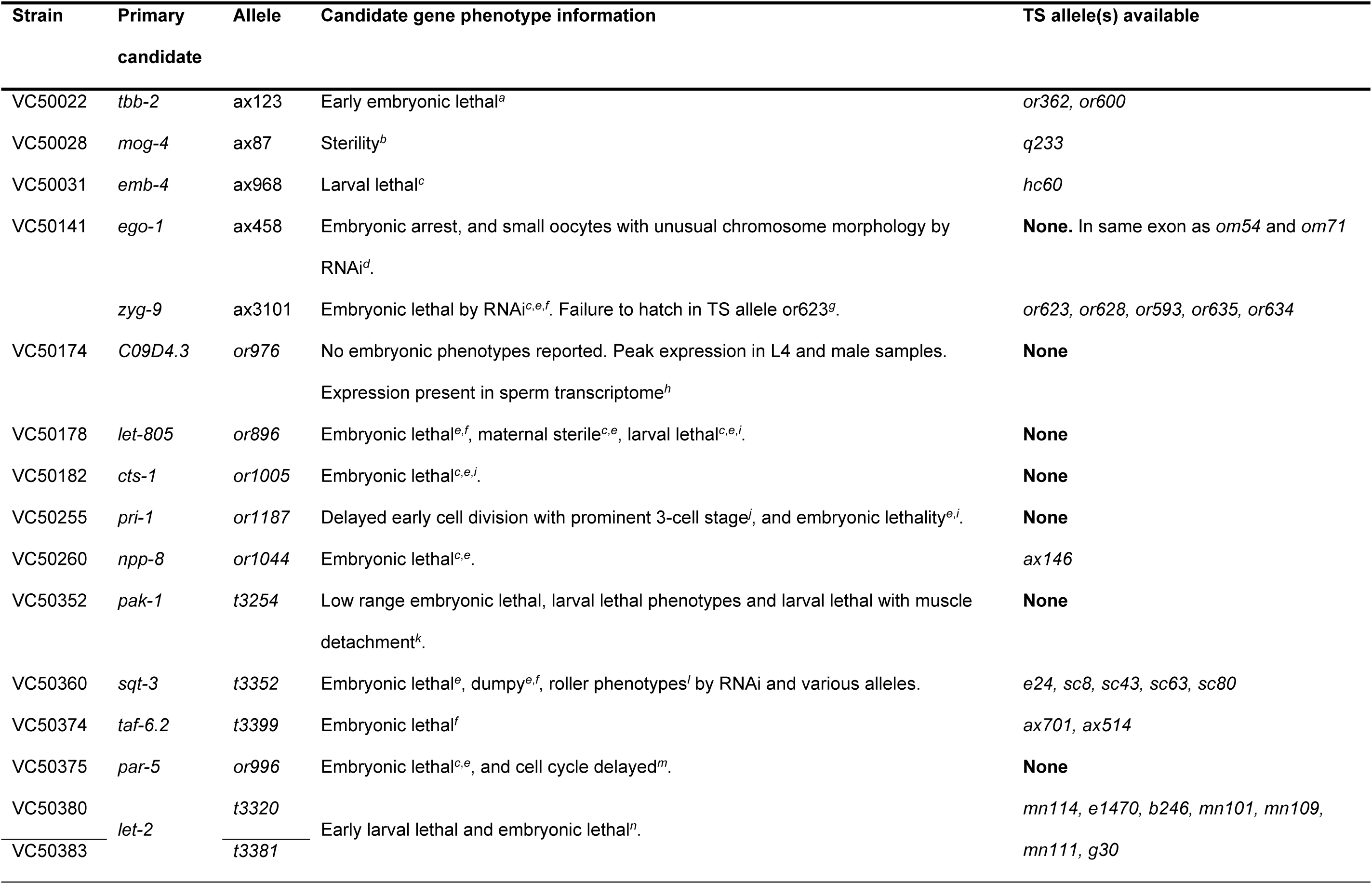
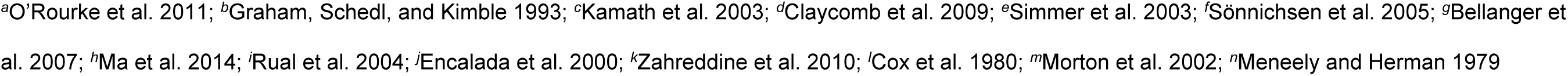
TS Candidate mutation information

## Discussion

Our large collection of sequenced *C.elegans* conditional mutant strains offers a rare opportunity to study essential genes on a larger scale. A high-throughput mapping of the causative alleles in this collection, however, is not a trivial task. We began our analysis of these strains with the goal to generate a robust mapping strategy that was affordable, high-throughput, comparable in resolution to current mapping paradigms and flexible enough for the community’s needs. Using single-molecule molecular inversion probes we developed the MIP-MAP strategy which relies on targeted loci sequencing of the *C.elegans* genome at great depth. MIP-MAP is differentiated from other mapping methods by 1) its adaptability to different mapping and segregant selection schema, 2) its independence from the usage of the Hawaiian strain CB4856, 3) by the reduction in sequencing and data generation while maintaining 1Mb resolution, and 4) its unique advantage in mapping the more recent collections of sequenced strains.

MIP-MAP successfully identified a phenotype-associated locus in the direct-selection of *sma-9* F2 segregants – a common mapping paradigm. However, with the *hlh-1(cc561)* temperature sensitive lethal allele, we successfully used a competitive selection scheme to identify the *hlh-1* locus versus an alternative approach such as picking dead larvae for WGS (Smith *et al.* 2016). This indirect approach to identifying lethal loci may also useful in gauging the rate of fixation across multiple conditions to elucidate mutation penetrance or gene function. In fact, our *hlh-1* data suggested that even at the permissive temperature of 16°, *hlh-1(cc561)* animals have some level of decreased fitness, despite a lack of reported lethality in this condition (Harfe *et al.* 1998).

Our results demonstrated the flexibility of this method to work with non-standard mapping strains. We leveraged a Million Mutation Project strain with a mere 269 SNVs and targeted strain-specific markers at 89 loci, using them to successfully map *sma-9* and *hlh-1* mutant alleles to within ~2Mb intervals. Furthermore, we have shown with VC20019 that the Million Mutation Project is also a library that contains polymorphic mapping strains with fewer potential issues than more divergent genomes like the Hawaiian strain. Based on our results it is even plausible that one could identify Million Mutation Project strains compatible for mapping synthetic phenotypes. This strategy would reduce the additional steps required in homozygosing background mutations used in forward genetic screens – simplifying the cloning process.

In terms of cost, the MIPs themselves are a “one-time” purchase where a single order at 25nM scale provides more than 2 million capture reactions per probe. MIP-MAP also offers comparable resolution to WGS with far less extraneous sequencing data. For example, each MIP-MAP library can be sequenced to an average 200,000 reads to obtain mapping data. In contrast, whole genome sequencing at recent standards of ~20X coverage (50bp, paired end) requires approximately 20 million reads - 2 additional orders of magnitude with most of the sequence being reference genome data. It is preferable to heavily multiplex samples during the sequencing process and thus lower overall cost per sample. Choosing the appropriate mapping strategy (bulk segregant versus competitive fitness) one could easily MIP-MAP fifty to one hundred times the number of strains on a single HiSeq lane compared to conventional whole genome sequencing mapping. The overall cost savings per strain and capabilities of MIP-MAP are therefore suitable to a high-throughput analysis of collections such as our TS strains or the Million Mutation Project. More recent large-scale genome wide association studies may also benefit from this method when verifying hits from amongst multiple candidate associations (Cook *et al.* 2016).

Although the MIP-MAP method was developed with sequenced mutant collections in mind, it should be noted that it is also an advantageous companion approach to standard genome sequencing and mapping of *de novo* strains. Million Mutation Project findings suggest genome sequencing to a 12-fold depth (Thompson *et al.* 2013) is sufficient to confidently identify SNVs and other mutations in a heavily mutagenized background. In combination with MIP-MAP, this still represents a potential sequencing savings of more than 7 million reads. Alternatively, once a mutation is mapped to a particular locus, the genomic interval pull-down sequencing method (O’Rourke *et al.* 2011b) can target chromosomal regions using *C. elegans* genomic Fosmids. These isolated intervals can be sequenced to identify mutations for that specific genomic interval. Thus, the throughput scalability of MIP-MAP still provides a significant advantage in mapping novel strains.

Intriguingly, a number of our mappings identified the presence of additional non-TS loci that conferred lowered fitness rates on the TS strains, suggesting that our TS collection holds a diverse library of additional mutations worth cloning. In a similar fashion, MIP-MAP could be useful in high-throughput mining of the Million Mutation Project for specific phenotypes in a form of reverse-genetics screen. For instance, rather than testing for temperature-sensitive lethality, MMP strains could be queried for phenotypes related to food source, plate conditions, or RNAi interactions. More broadly, MIP-MAP demonstrates, on a focused level, a method of barcoding strains within the context of a mixed population. This low-cost high-throughput form of analysis could be applied, for instance, to the analysis of mixed pools of multiple strains to track variant fitness across a diverse set of conditions. Targeting strains in this manner might reveal the functions of many novel genes. Given that the Million Mutation Project strains have already been sequenced, using MIP-MAP to identify the cause of these phenotypes would generate far less sequencing data than WGS methods.

Reviewing our TS-mapping workflow, five of nine strains were bulk segregant mapped with more than 60 F2s chosen as TS-positive but none of these generated a concise interval – rather they complemented the sharp intervals obtained by the competitive fitness approach. Curiously, in three of these five sets the chosen F2 population was far less than the expected 25% ratio for a homozygous recessive mutation (**Supplemental Table S1**). Indeed, in the remaining four strains mapped with F2 counts below 60, these animals were identified at low frequency from very large populations of potential F2s – further suggesting issues in either identifying the TS phenotype or reduced phenotype penetrance in these populations. In *toto*, using bulk segregant analysis we observed only three strains (VC50028, VC50374, and VC50380) that generated mapping intervals comparable in resolution to data from the competitive fitness approach. In that respect, a MIP-MAP competitive fitness assay over many generations offers the ability to identify mutant loci that are of incomplete penetrance and may prove problematic in standard bulk segregant mapping. From this we conclude that for the remainder of our TS strains – especially those of apparently lower penetrance – it is more ideal to complete both mapping approaches in parallel with the benefit of reducing repetition of mapping crosses in downstream workflows.

The collection of *C. elegans* TS strains we present here is, to our knowledge, the largest, fully-sequenced collection to date. Based on our mapping results we suspect that many new and interesting alleles can be mined from these strains. For example, our mapping of the strain VC50174 identified a sperm-specific essential gene *C09D4.3*. Recent studies suggested that sperm-targeted RNAi screens may be inefficient due to competition between endogenous and exogenous RNAi pathways. This hypothesis supports the literature from multiple RNAi screens that report no embryonic lethal phenotypes for *C09D4.3* (Fraser 2000; Maeda *et al.* 2001; Sönnichsen *et al.* 2005). Finding this TS allele within our small set suggests that our overall collection of strains has the potential to not only yield the first TS alleles for a number of essential genes but also the potential to identify new essential genes not discovered by standard RNAi screens. Further analysis of our collection as a whole may identify that genes with a high frequency of mutant alleles in TS loci could help to prioritize candidate intervals. Our data also suggests that ~6% of the strains in this collection may carry TS alleles for genes not previously characterized as essential.

In summary, we present a large collection of sequenced temperature sensitive mutants housing a variety of embryonic-lethal and sterility phenotypes. In conjunction with this, we demonstrate the MIP-MAP methodology as a robust, cost-efficient, and high-throughput genetic mapping format. It allows for a wide range of flexible sample generation schema that can be implemented with relative ease. We believe MIP-MAP can even identify alleles with weak penetrance phenotypes given the proper selection conditions. Taken together, these two tools present a unique opportunity for the nematode research community to investigate a diverse library of mutant strains and to map these and other mutant alleles in a unique and accessible manner. We believe both of these resources will be a beneficial addition to the nematode and scientific community at large.

## Acknowledgments

We thank Emily Turner, Joseph Hiatt and Choli Li for advice on MIP library preparation and sequencing. C.A.M is supported by the Canadian Institutes for Health Research MFE-135408. N.M is supported by the Internationales Graduierten Kolleg Niedersachsen. Work in the B.B. laboratory was supported by an R01 grant GM114053 from the NIH. Work in the G.S. laboratory was supported by an R01 grant HD37047 from the NIH. G.S. is an Investigator of the Howard Hughes Medical Institute. Work in the D.G.M laboratory was supported by the Canadian Institutes for Health Research and the NHGRI (through R.H.W). D.G.M is a Senior Fellow of the Canadian Institute for Advanced Research. Work in the R.H.W laboratory was supported by an ARRA GO grant HG005921 from the NHGRI, an R21 grant HG007201-02 from the NIH, and by the William H. Gates Chair of Biomedical Sciences.

---

**Supplemental Figure S1.**
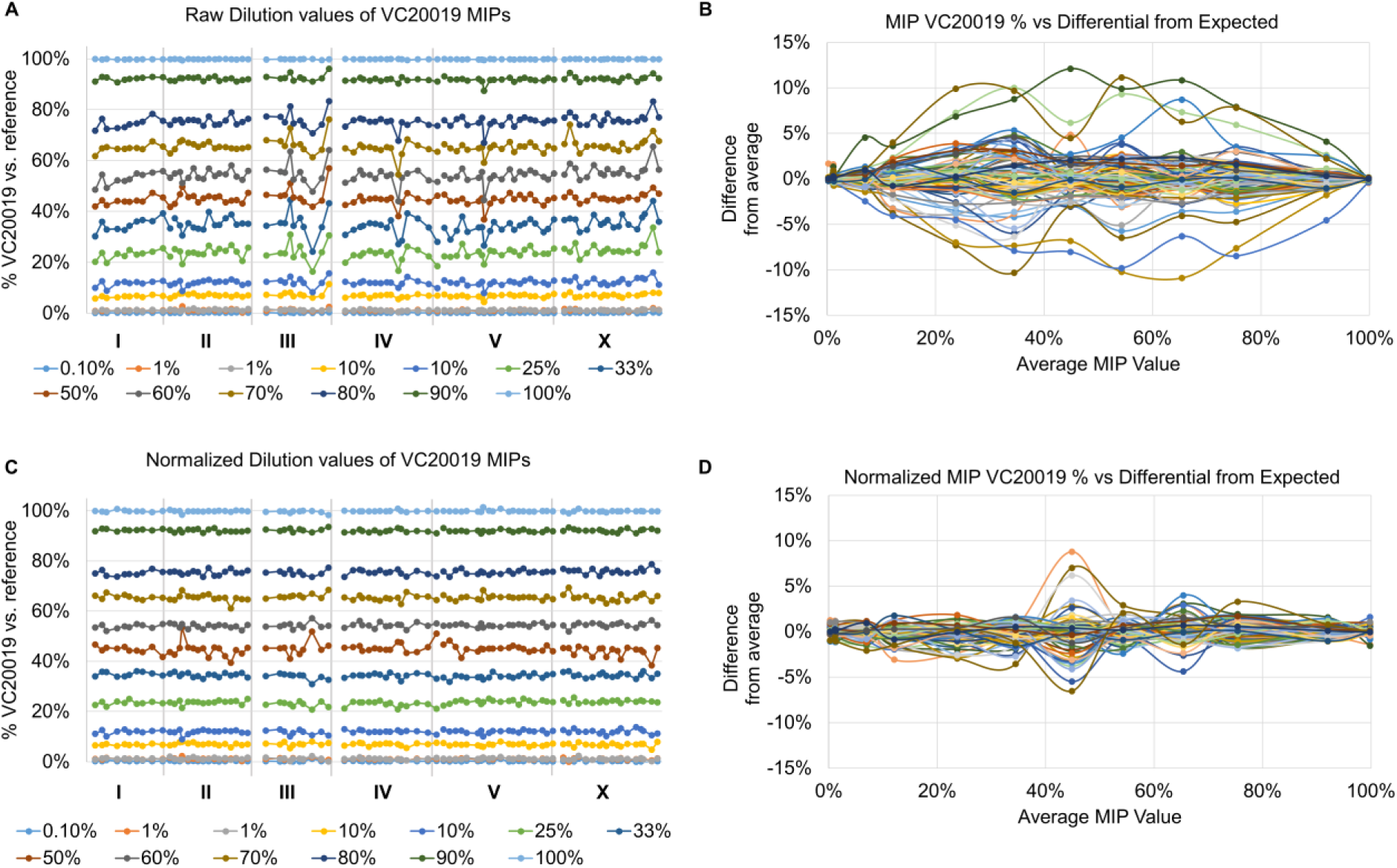
MIPs generate consistent results but may require normalization. The concordance of SNV ratios across multiple loci and sample dilutions (A), identified a small subset of MIPs that generated incorrect read calls across multiple dilutions. The difference between the called dilution value and the average across all 90 MIPs was used to generate a behavioural curve (B). Behavioural curves were used to normalize the MIP SNV ratios, generating more consistent calls across all MIP targets (C, D).

**Supplemental Figure S2.**
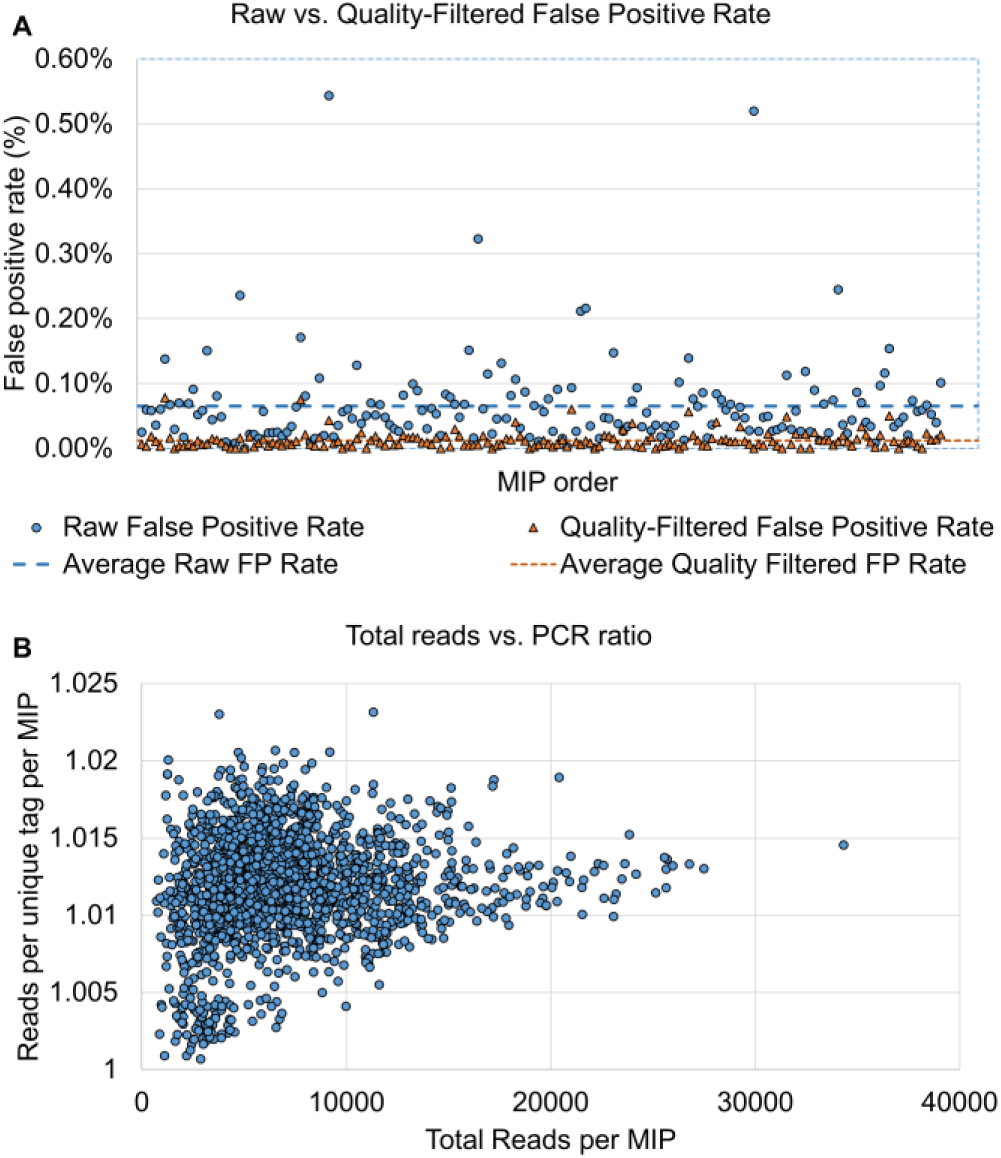
MIP accuracy and efficiency were tested on a range of samples. (A) The false positive rates of MIPs were investigated using a series of probes targeting 172 genomic loci. A mean raw false positive read rate of 0.0651% ± 0.0696% was observed with a maximum of 0.543%. By filtering reads at the SNV site for quality scores of Phred >= 30, the false positive rate was decreased to a mean of 0.0122% ± 0.0129% with a maximum value of 0.0785%. (B) MIP capture and library preparation produces a high concordance of sequence reads vs. molecular tags. Using 24 sets of MIP capture reactions with the same 89 MIPs, the observed mean rate of molecular tag duplication via PCR is 1.048 ± 0.006. The ratio of reads per tag was not correlated to the total reads per MIP when analysing the same 24 sets by total read count on a per MIP per capture reaction basis.

**Supplemental Figure S3.**
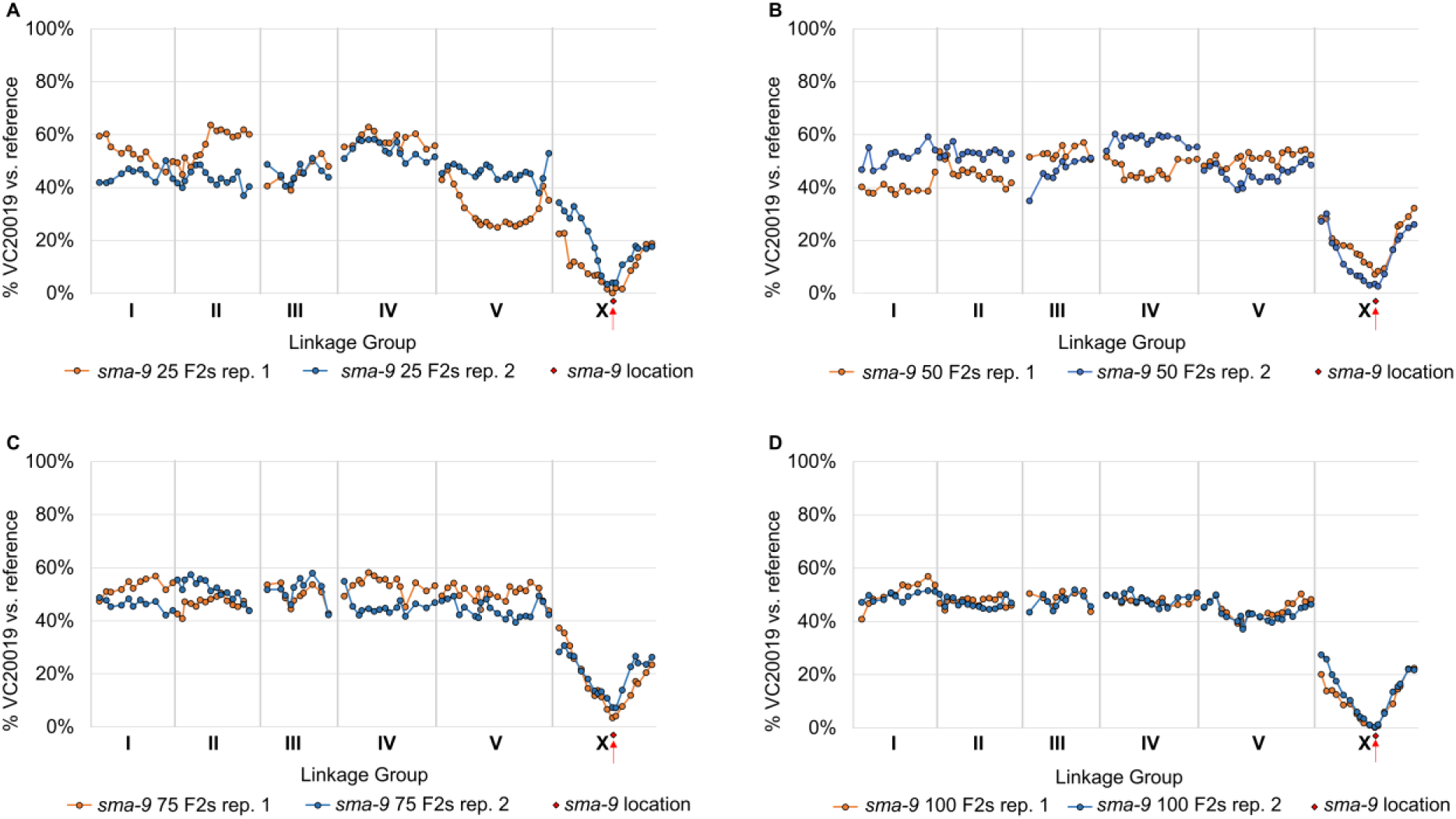
MIP-MAP profiles for *sma-9* mutants improves when using more F2 recombinants. A sufficient MIP-MAP profile to identify the *sma-9* locus was generated with as few as 25 F2 *sma-9* phenotype animals (A) but mapping improved with increasing numbers of F2s populations at 50 (B), 75 (C) and 100 (D) F2 animals.

**Supplemental Figure S4.**
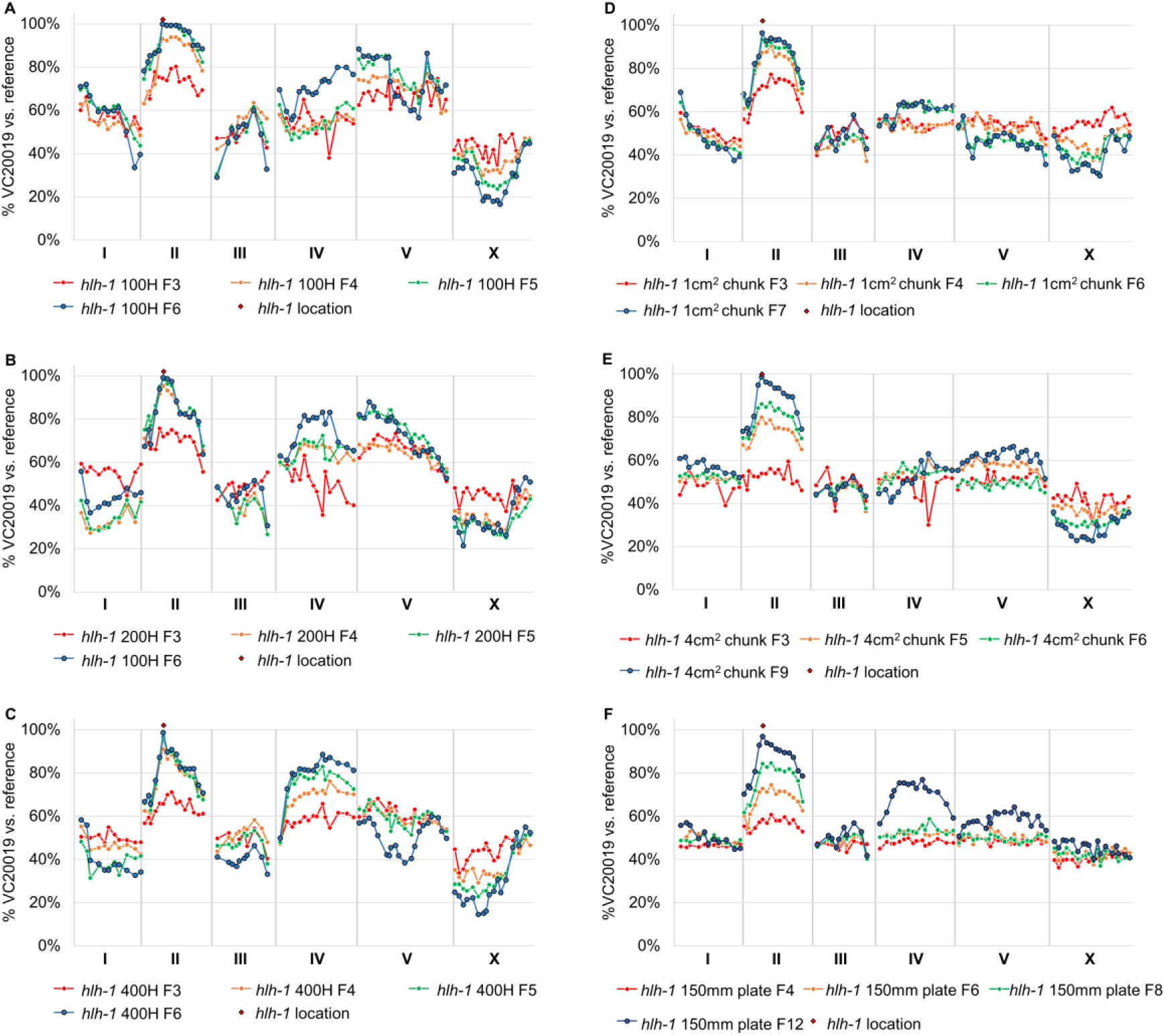
MIP-MAP profiles for *hlh-1* mutants can be generated by indirectly mapping in multiple schemes. MIP-MAP mapping profiles for *hlh-1* were reproduced using varying numbers of animals transferred each generation until fixation against the TS *hlh-1(cc561)* allele occurred. The size of population transferred each generation was tested at 100 (A), 200 (B), and 400 (C) animals. Random selection of animals was verified populations were transferred by chunking 1cm^2^ (D) or 4cm^2^ (E) of animals or washing the entire population and portioning ~10,000 animals onto new 150mm plates (F). Each line present in a graph represents mapping data from a different generation for the same experimental replicate.

**Supplemental Figure S5.**
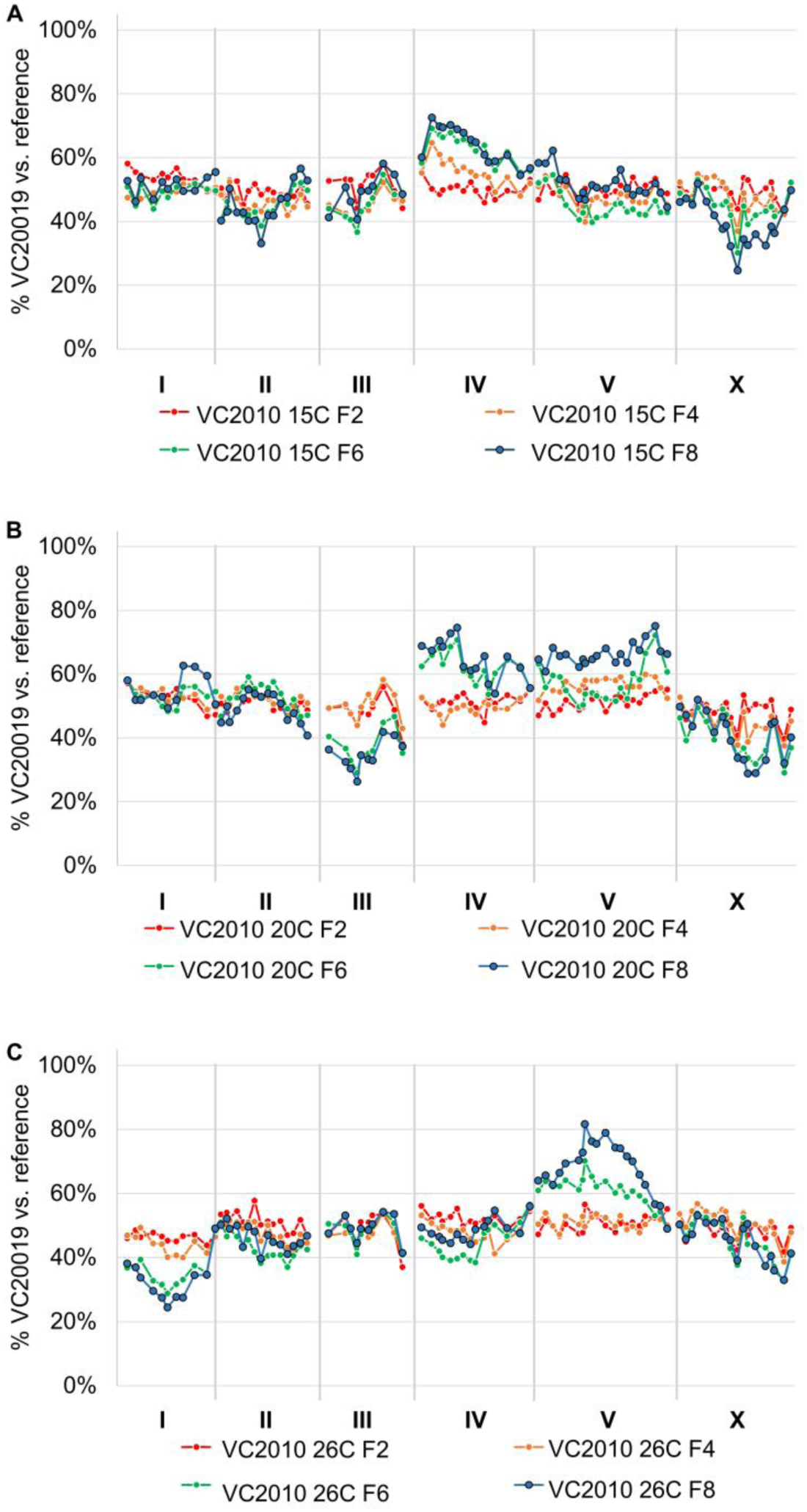
VC20019 appears to have little to no temperature-sensitivity. The mapping profile of a cross between VC20019 and VC2010 was completed with very little fixation at any particular locus across the genome after mapping at 15°C (A), 20°C (B), and 25°C (C) for at least 8 generations. This data indicates that the mapping mutant VC20019 has comparable fitness to our N2 reference *C. elegans* strain VC2010.

**Supplemental Figure S6.**
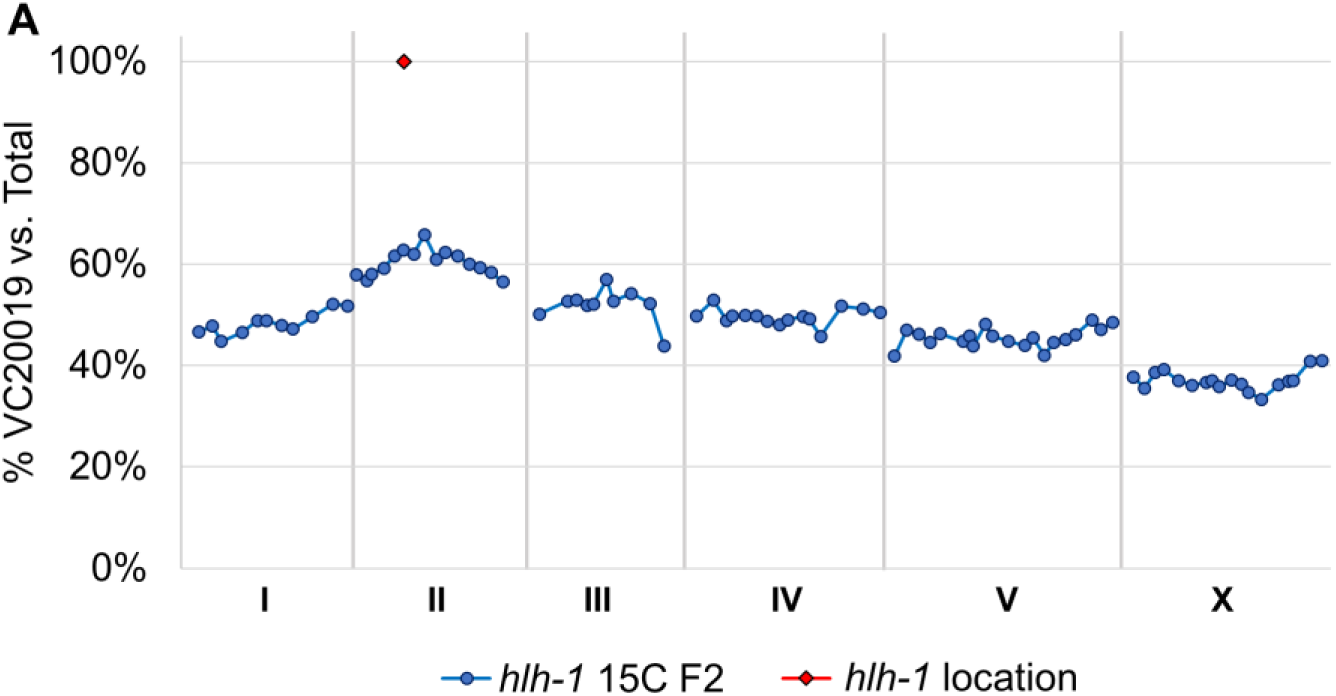
The *hlh-1(cc561)* allele exhibits mild fitness defects at a permissive temperature. While mapping the *hlh-1(cc561)* allele at 15°C the beginnings of fixation against the *hlh-1* interval was observed, even after a single generation at the permissive temperature.

**Supplemental Figure S7.**
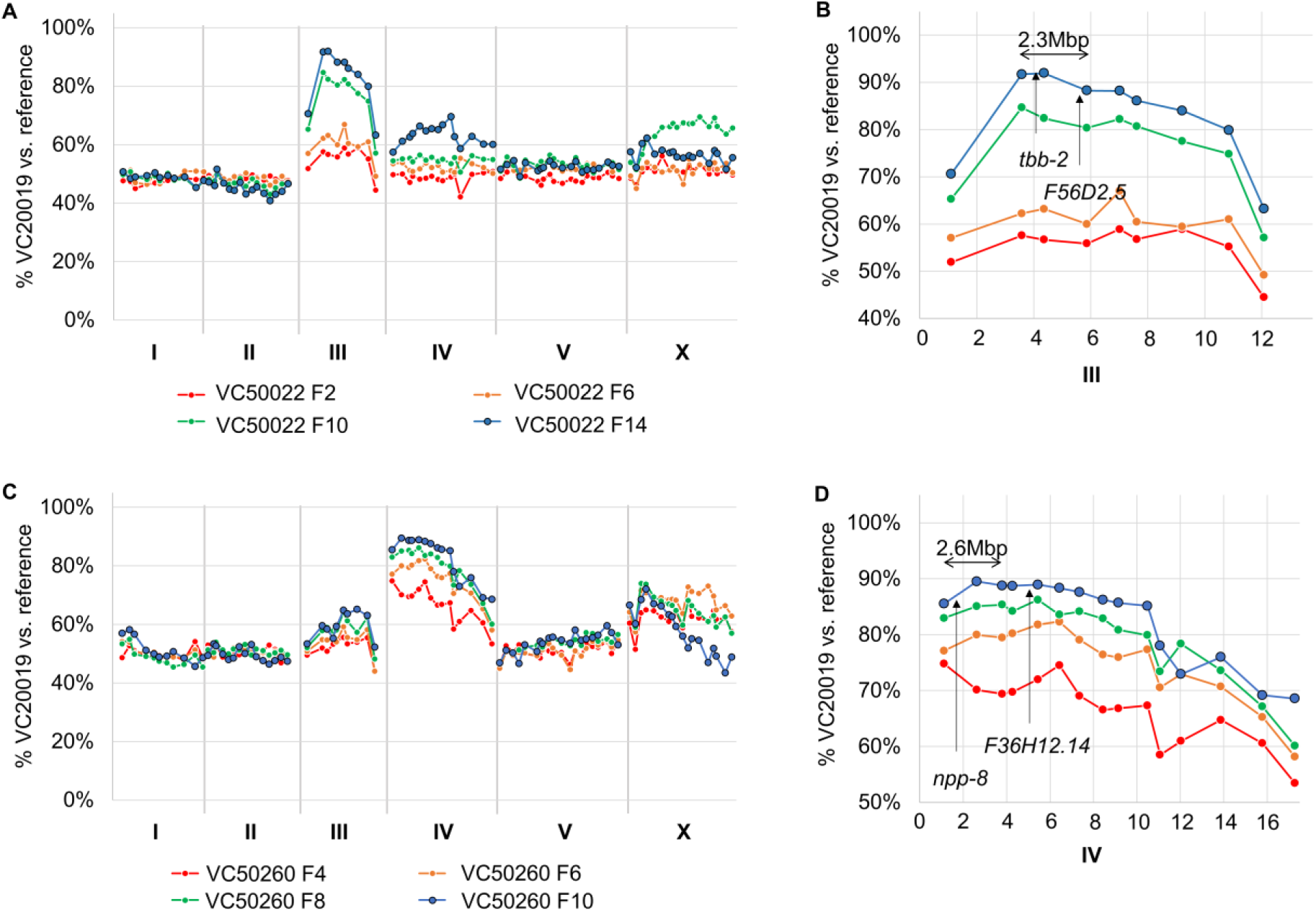
TS strains that resolved with a single locus. VC50022 is an example of a clean TS mapping that identified a single locus without additional loci (A). In this strain there are only 2 relevant coding mutations within the identified interval on LGIII (B). The most likely candidate *tbb-2* has a number of embryonic lethal phenotypes. VC50260 (C) resolved without much indication of any other candidate loci. Within this interval on LGIV there was only 1 relevant coding variant which was located in *npp-8* (D). The next closest relevant coding variant is in *F36H12.14* nearly 1.5Mbp from the predicted interval.

**Supplemental Figure S8.**
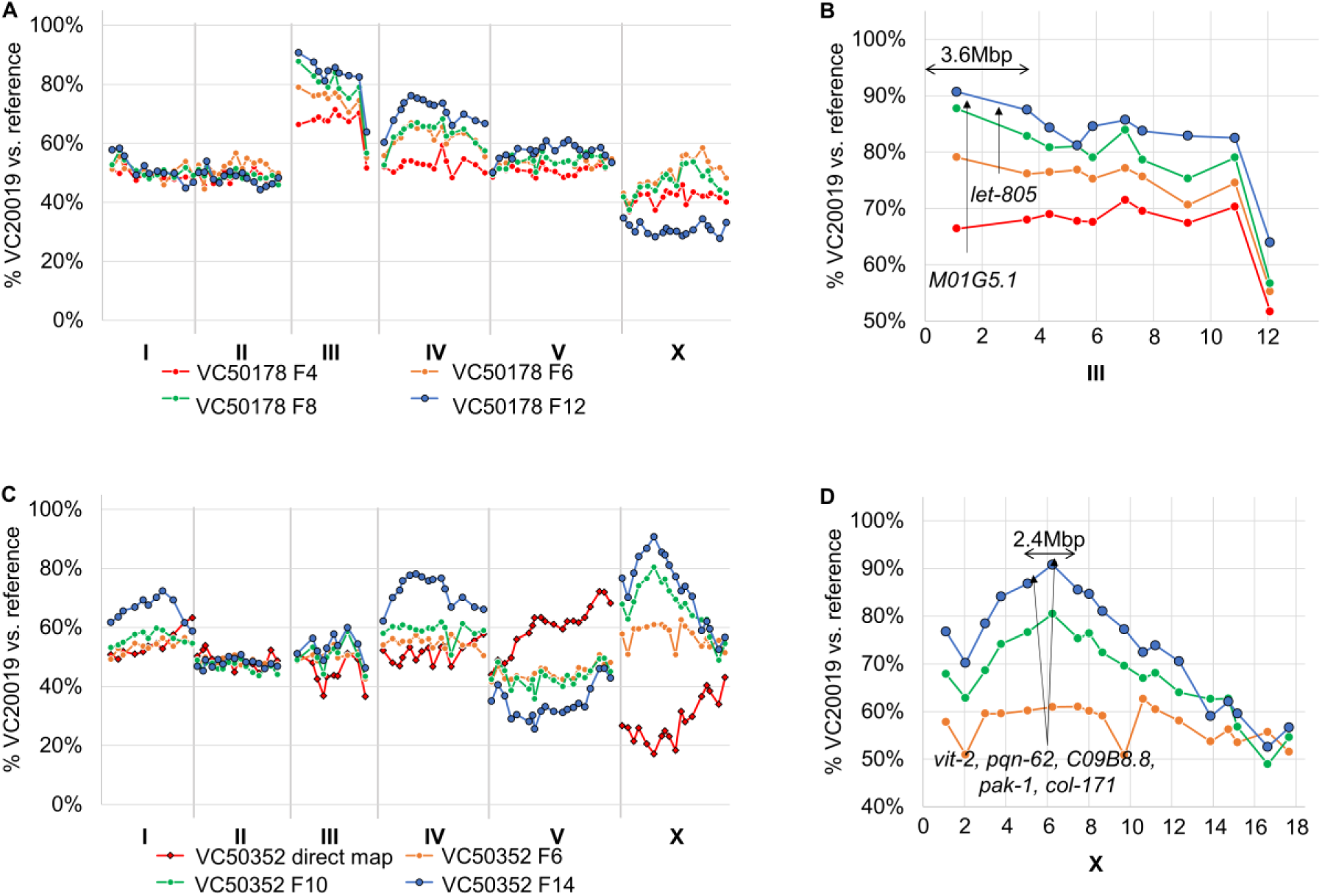
TS strains with mapping profiles that identify additional weak alleles of low fitness. The TS locus within VC50178 (A) was identified on the left arm of LGIII by following competitive fitness mapping over multiple generations. The predicted 3.6Mbp interval (B) has only two relevant coding changes with the likely candidate as *let-805* based on phenotyping of the VC50178 strain. The TS locus on LGX in VC50352 (C) was resolved with the liquid bulk segregant mapping assay. Within this interval (D) are 7 relevant candidates with the prime candidate being in *pak-1* based on the VC50178 observed TS phenotype. It is possible the additional loci harbour weak or low-penetrance alleles involved in population fitness and are not apparent until the TS-allele has been removed from the population.

**Supplemental Figure S9.**
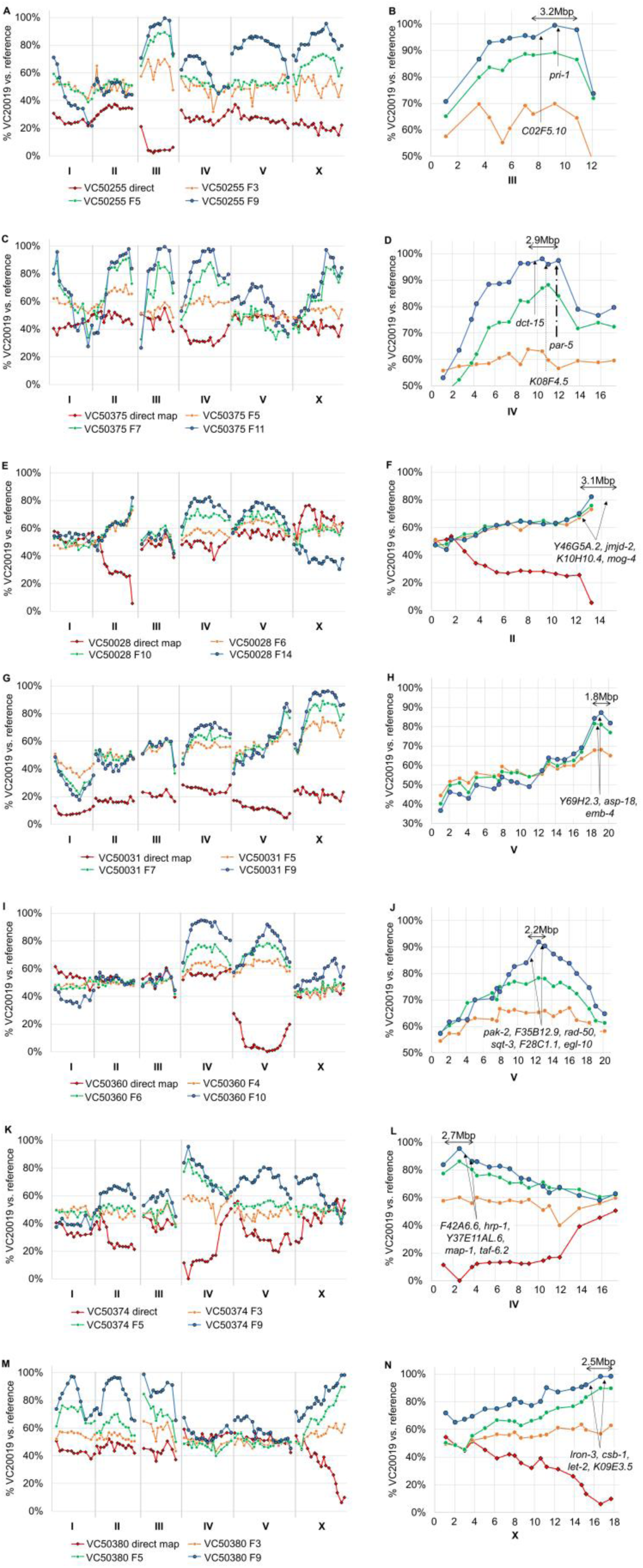
TS strains can carry multiple strong fitness-defective loci. In three cases, the competitive fitness mapping protocol identified multiple loci within these strains that conferred an overall reduced fitness to the population at a fixation rate that made it difficult to identify the correct TS-specific locus. Additional mapping with the liquid-format bulk segregant assay elucidated the correct linkage group for the TS phenotype - albeit with lower resolution. The TS-associated locus in VC50255 was identified at LGIII (A) with the prime candidate as *pri-1* based on phenotype and Wormbase reports (B). A 2.9Mbp TS interval was identified in VC50375 on LGIV (C) containing 3 candidates (D) with the prime candidate as *par-5* based on a shared cell-cycle delay phenotype in the strain. In VC50028 a 3.1Mbp interval on LGII (E) was identified with *mog-4* as the primary candidate based on shared phenotype data between the strain and Wormbase sources (F). The strain VC50031 had multiple loci with a TS-associated hit on a 1.8Mbp interval on LGV and a likely hit for *lin-2* on LGX (G). The primary candidate in this interval is *emb-4* (H) based on based on VC50031’s sterility phenotype and a reported high early embryonic expression of this gene. The strain VC50360 produced a mapping profile (I) with two very strong candidate loci that were resolved to a 2.2Mbp interval on LGV (J) with a primary candidate of *sqt-3* as the strain’s phenotype overlapped with reported RNAi and allele phenotypes. VC50374 exhibited a number of peaks that were mirrored in liquid-format segregant mapping with the TS-locus resolved at LGIV (K). The 2.7Mbp interval (L) contained many candidates with the primary candidate as *taf-6.2*, a gene reported to have embryonic lethal phenotypes and TS lethal alleles. One suc allele is located in the same exon as the VC50374 SNV. The TS strain VC50380 (M) identified 4 strong low-fitness loci with liquid-format bulk segregant mapping resolving the TS locus to LGX. Within this interval (N) the primary candidate is *let-2* based on reported phenotypes compared to this strain.

**Supplemental Table S1.** 96-well liquid mapping number of F2 segregants used

| Strain | Set | Direct Mapping number of<br>F2s (chosen/total) | % F2s Picked | Mapping observations |
| --- | --- | --- | --- | --- |
| VC50352 | 2 | 37/362 | 10.22 | Clear peak by F10 but carried to F14. Direct mapping locus only fixates to 20%. |
| VC50255 | 1 | 66/440 | 15.00 | Clearest interval by F9. Direct mapping has very broad interval across most of<br>LGIII. |
| VC50375 | 1 | 109/453 | 24.06 | Clear peak at F7 but also matched in amplitude by 4 other peaks. Direct map<br>resolved chromosome but with poor resolution. |
| VC50028 | 3 | No Data | No Data | Quick fixation observed on LGII to 80% by F6 but does not improve even at F15.<br>Direct mapping mirrored exact interval. |
| VC50031 | 3 | 72/737 | 9.76 | Clear interval on LGV by F7 but best at F9. Direct mapping has high signal noise<br>but confirms LGV. |
| VC50141 | 3 | 75/387 | 19.37 | Mapping doesn't clarify until between F8 and F12. Direct mapping fixates to<br>~30% on each locus. |
| VC50360 | 2 | 98/439 | 22.32 | Clearest interval on LGV by F10. Direct mapping confirms but with a very broad<br>interval across most of LGV. |
| VC50374 | 2 | 15/384 | 3.90 | Clear interval on LGIV by F5 but best fixation by F9. Direct mapping managed to<br>identify the interval with 15 F2s. |
| VC50380 | 2 | 42/384 | 10.93 | Somewhat clear peak on LGX by F5 but reached fixation at F9 with 3 additional<br>strong intervals. Direct mapping confirmed LGX with good resolution. |

- CF = competitive fitness mapping - 96-well = liquid bulk segregant mapping

